# Fast and accurate estimation of selection coefficients and allele histories from ancient and modern DNA

**DOI:** 10.1101/2023.12.16.572012

**Authors:** Andrew H. Vaughn, Rasmus Nielsen

## Abstract

We here present CLUES2, a full-likelihood method to infer natural selection from sequence data that is an extension of the method CLUES. We make several substantial improvements to the CLUES method that greatly increases both its applicability and its speed. We add the ability to use ARGs on ancient data as emissions to the underlying HMM, which enables CLUES2 to use both temporal and linkage information to make estimates of selection coefficients. We also fully implement the ability to estimate distinct selection coefficients in different epochs, which allows for the analysis of changes in selective pressures through time. In addition, we greatly increase the computational efficiency of CLUES2 over CLUES using several approximations to the forward-backward algorithms and develop a new way to reconstruct historic allele frequencies by integrating over the uncertainty in the estimation of the selection coefficients. We illustrate the accuracy of CLUES2 through extensive simulations and validate the importance sampling framework for integrating over the uncertainty in the inference of gene trees. We also show that CLUES2 is well-calibrated by showing that under the null hypothesis, the distribution of log-likelihood ratios follows a chi-squared distribution with the appropriate degrees of freedom. We run CLUES2 on a set of recently published ancient human data from Western Eurasia and test for evidence of changing selection coefficients through time. We find significant evidence of changing selective pressures in several genes correlated with the introduction of agriculture to Europe and the ensuing dietary and demographic shifts of that time. In particular, our analysis supports previous hypotheses of strong selection on lactase persistence during periods of ancient famines and attenuated selection in more modern periods.

## Introduction

One of the primary evolutionary forces that shapes the genetic variation of populations is natural selection, the causal effect of genotype on the reproductive success of an individual. While experimental evolution studies can enable a somewhat direct measurement of natural selection in certain limited cases, most methods for inferring natural selection rely on statistical frameworks to analyze a sample of sequence data. Given that sequence data lies in a very high-dimensional space, these methods for inferring natural selection have historically been based on summary statistics, that can be thought of as projections of the sequence data to a lower-dimensional subspace. These summary statistics include SNP-frequency statistics such as Tajima’s D (Tajima, 1989) or Fay and Wu’s H (Fay and Wu, 2000), as well as linkage disequilibium statistics such as the extended haplotype homozygosity statistic (Sabeti et al., 2002) or the LD decay test (Wang et al., 2006). One can compute these statistics for a given sequence, or in sliding windows across the genome, and conclude that values significantly different from their null expectation are indicative of natural selection. As an extension of the basic summary statistic approach, approximate Bayesian computation (ABC) frameworks, which can use information from many different summary statistics, have also been used to infer selection coefficients (Peter et al., 2012). A variety of machine learning methods to detect selection have additionally been developed (Schrider and Kern, 2018; Torada et al., 2019; Hejase et al., 2021), which all rely on training models on a set of features obtained from sequence data. However, these approaches ignore a large amount of the information present in the data by only considering sets of summary statistics or features. Furthermore, these methods lack flexibility, for example if one wishes to specify different parameters or classes of models, such as allowing for selection coefficients to differ in different time periods or incorporating ancient samples that are sampled at distinct time points.

Overall, therefore, a computationally tractable full-likelihood method is needed that does not rely on summary statistics, and can thus use all the information present in the data, and which also has the flexibility to be able to be applied to a variety of datasets and models. It is to this end that the method CLUES was developed by (Stern et al., 2019). The original CLUES method is based on the observation that if one can determine a full likelihood expression for the data conditional on the full history of the frequency of an allele, then one can consider the allele history as a latent variable and integrate over the full set of possible allele frequencies in order to obtain an expression for the full likelihood function. As the sequence data and selection coefficient of an allele are approximately independent conditional on the gene tree at that locus, the full likelihood of the data could be computed by computing the likelihood of that gene tree. In reality, the true gene tree at a site is never known with certainty but instead must be estimated. Estimating gene trees in recombining species is challenging as recombination causes neighboring regions of the genome to have distinct, but correlated, gene trees. The full collection of these correlated gene trees across a genomic region is called the *ancestral recombination graph* (ARG), the theory for which was first developed by (Hudson, 1983) and (Griffiths and Marjoram, 1996). The key insight of using ARGs for population genetic inference is that they contain all the information about the history of each region of the genome. Therefore methods that employ ARGS are utilizing the maximum possible information present in the data. The inference of ancestral recombination graphs is made difficult by the large state space of possible graph topologies and the relatively small amount of information mutations provide about the underlying graph. However, recent methods to infer ARGs have made significant progress on this difficult computational task (Rasmussen et al., 2014; Kelleher et al., 2019; Speidel et al., 2019; Mahmoudi et al., 2022; Zhang et al., 2023). Nevertheless, there will still always be uncertainty as to the underlying ARG, as there will be many ARGs that are compatible with the observed sequence data. For that reason, CLUES also integrates over the uncertainty in the ARG estimation, in addition to integrating over the set of possible allele histories, thus giving two sets of latent variables that are integrated out to obtain the full likelihood function.

We here present CLUES2, an extension of the CLUES method, which is able to work not only on importance samples of ARGs on modern data but also on ancient genotype samples and ARGs built on ancient data, thus enabling the usage of time series data and linked SNPs to give better estimates of the selection coefficient of an allele. We also develop and test the capability of CLUES2 to estimate different selection coefficients in different time periods. In addition, we make significant improvements to runtime by making several well-justified approximations to the forward and backward algorithms. The computational speedups allow more importance samples to be used, which increases accuracy and also allows CLUES2 to be used in genome-wide scans of selection, where hundreds of thousands or millions of SNPs may need to be analyzed. We perform rigorous testing of both selection coefficient estimation and the appropriateness of the chi-squared test for hypothesis testing of selection. Finally, we improve the interpretability of our reconstruction of historic allele frequencies by integrating over the uncertainty in the estimation of the selection coefficients. We make CLUES2 available on GitHub as a well-documented Python package at https://github.com/avaughn271/CLUES2.

## Materials and Methods

### CLUES2 Framework

We here describe the basic hidden Markov model framework of CLUES2. We wish to find the value of *s*, the selection coefficient of the derived allele, that maximizes the likelihood of the data *D*. What exactly *D* represents depends on the kind of data being analyzed, and we postpone a formal definition of *D* until the subsequent sections. We compute this likelihood by conditioning on the historic trajectory of the derived allele frequency. In particular, if we let *χ* be the set of all possible derived allele trajectories, we compute the likelihood as:

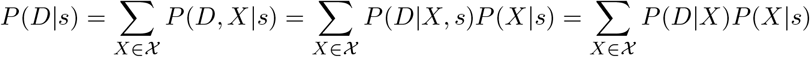

where the last equality follows from the fact that the data is independent of the selection coefficient given the allele trajectory. If we discretize the derived allele frequency into *K* frequency bins and consider the history of an allele until a maximum time point *T* (measured in discrete generations), computing this expression would naively require summing over *K*^*T*^ many allele trajectories. However, we instead recognize this as a hidden Markov model (HMM) where the derived allele frequency at a given time is the hidden state and the data are the emissions. We can then efficiently compute this sum using the forward algorithm. This modeling of allele frequencies as hidden states of an HMM has previously been applied by (Williamson and Slatkin, 1999), (Bollback et al., 2008), (Steinrücken et al., 2014), (Bergman et al., 2018), and (Paris et al., 2019).

The implementation of the forward algorithm is as follows: we compute a forward algorithm matrix *F*, which has dimension *K* × *T*. The first column is initialized to all zeros, except that a 1 is placed in the row corresponding to the allele frequency bin closest to the observed modern allele frequency. Assuming the first *t* − 1 columns of *F* have been computed, one time-step of the forward algorithm consists of computing, for each frequency bin *k*,

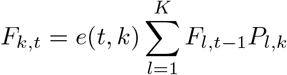

where *P*_*l,k*_ is the probability of transitioning from allele frequency bin *l* to bin *k* in one generation and *e*(*t, k*) is the probability of observing the data at time *t* given you are in state *k*. Given *K* allele frequency bins, we consider the *K* numbers that are equally spaced between 1*/*200000 and 1 − 1*/*200000 inclusive, call them *w*_1_, …, *w*_*K*_. We set the allele frequency of bin *k*, call it *x*_*k*_, to be the quantile function of a Beta(1/2,1/2) distribution at *w*_*k*_. This creates a spacing of numbers between 0 and 1 that is denser near the boundaries. If 1 *< l < K*, then *P*_*k,l*_ is approximated as Φ_*s,N*_ ((*x*_*l*_ + *x*_*l*+1_)*/*2) − Φ_*s,N*_ ((*x*_*l*−1_ + *x*_*l*_)*/*2) where Φ_*s,N*_ (*x*) is the CDF of a normal distribution with mean *x* − *sx*(1 − *x*) and variance 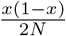. *P*_*k*,1_ is Φ_*s,N*_ ((*x*_1_ + *x*_2_)*/*2) and *P*_*k,K*_ = 1 − Φ_*s,N*_ ((*x*_*K*−1_ + *x*_*K*_)*/*2). We note that while this definition restricts population size *N* to be constant through time, we do indeed allow population sizes to vary through time (see section “Differing population sizes” for details). Furthermore, it is worth highlighting that throughout this manuscript, we use *N* to refer to haploid effective population size, denoting the number of sequences present in a population, not the number of diploid individuals in that population. After computing each entry of the matrix *F*, we have uniform exit probabilities from the chain, meaning that 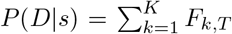. We then optimize *P* (*D*|*s*) with respect to *s* to achieve our estimate *ŝ*^*MLE*^. Dividing by *P* (*D*|*s* = 0) yields the log-likelihood ratio, which can then be used for hypothesis testing (see the section “Validation of chi-squared test”). In the following sections we discuss the types of data we use for the emissions *e*(*t, k*).

### Ancient Genotypes

One type of emissions considered are ancient genotypes. This is a list of genotypes of ancient individuals along with the times at which these individuals were sampled. If we denote a homozygous derived genotype as *DD*, a homozygous ancestral genotype as *AA* and a heterozygous genotype as *AD* we have

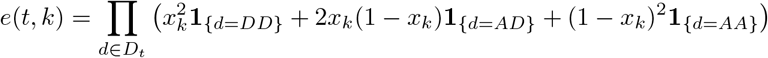

where *D*_*t*_ represents the set of all genotypes sampled at time *t*.

In practice, we allow the usage of genotype likelihoods as can be generated by the imputation of ancient genomes, so our expression becomes

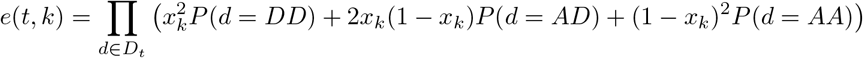

We validate this approach on simulated data (see Figure 1a). If our implementation is correct, there should be exactly two sources of error, which both contribute to variance of our estimator:

**Fig 1:**
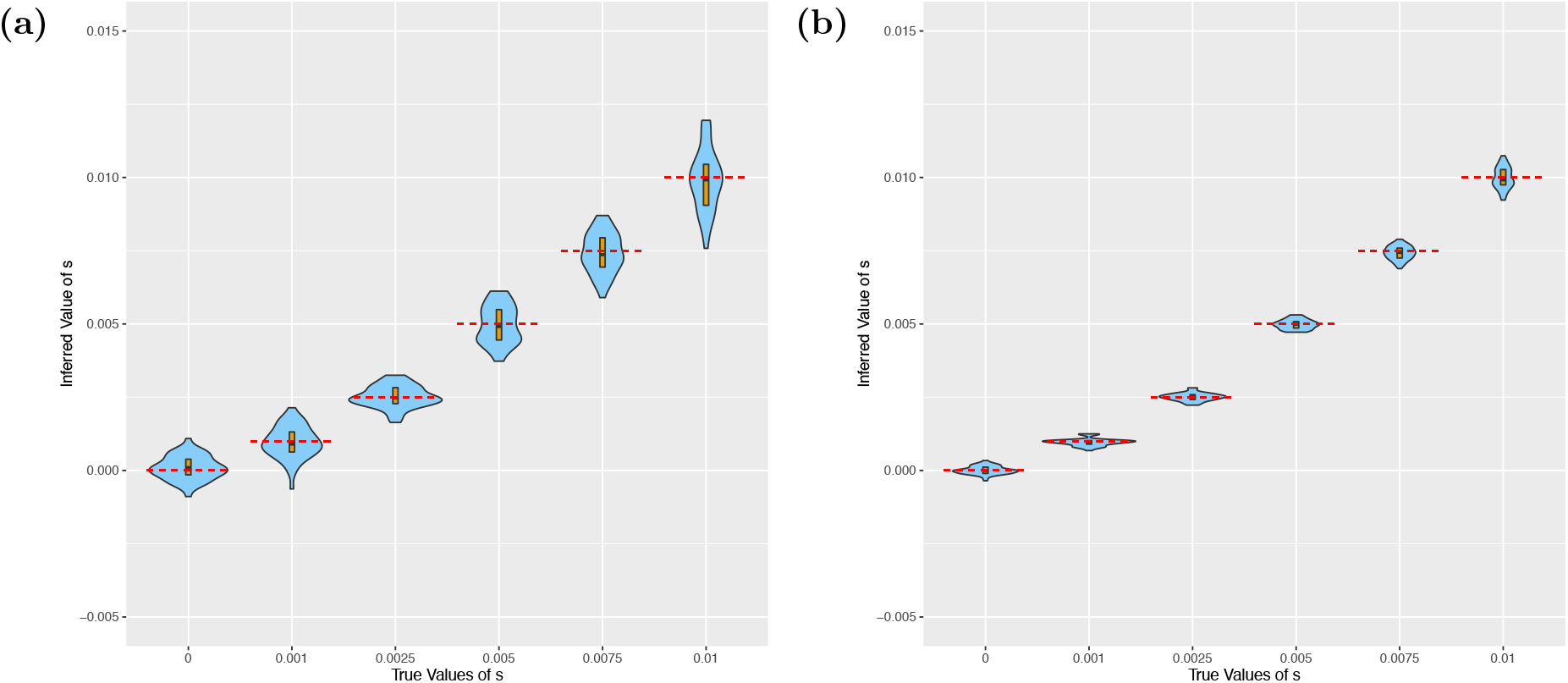
Violin plots showing the results of running CLUES2 on ancient genotype data. Boxplots are overlayed with the whiskers omitted. True values of *s* are shown as dashed red lines. 30 replicates were performed for each true value of *s*. Simulations were run with (a) *N* = 30000 and an individual sampled every 4 generations and with (b) *N* = 600000 and an individual sampled every 1 generation.

1. Fluctuations in the true allele frequency trajectory due to genetic drift, which causes the true allele frequency to be less informative of the true value of *s*
2. Noisy estimation of the true allele frequency due to finite sampling of ancient genotypes.

Therefore, we ran simulations with very low drift and very dense sampling of ancient genotypes (see Figure 1b). We observed that the estimated values of *ŝ*^*MLE*^ approximately converged to the true values, which validates the correctness of our implementation. We also allow for the usage of ancient haplotype probabilities, in addition to genotype probabilities, by substituting the expression *x*_*k*_*P* (*d* = *D*) + (1 − *x*_*k*_)*P* (*d* = *A*) in the product above.

### True Trees

The next class of emissions we consider are coalescence events of a gene tree. As in the original CLUES paper, we consider a structured coalescent model of the gene tree of a locus under selection following (Kaplan et al., 1988) and (Braverman et al., 1995). In particular, we assume we know the allele labeling at each leaf node of the tree and that the gene tree satisfies the infinite sites assumption with respect to the allele (we discuss violations to the infinite sites assumption later in the “Inferring Gene Trees on Ancient Human Data” section). We can then obtain a labeling of each branch in the tree as a derived allele branch or an ancestral branch and a labeling of each coalescence nodes in the tree as a derived allele coalescence or an ancestral branch (See Figure 2). The one exception to this is the branch on which the mutation must have arisen, which we call the mixed lineage.

**Fig 2:**
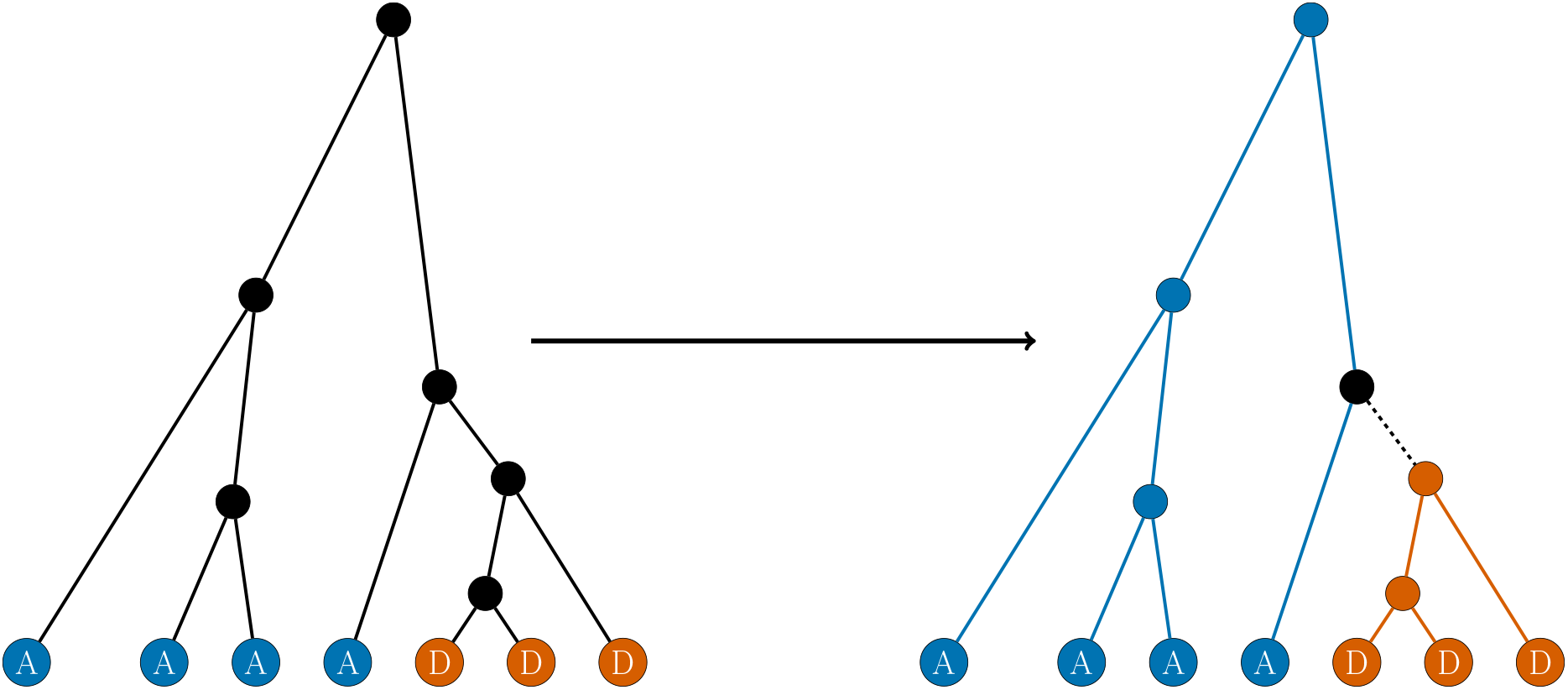
An outline of the labeling of branches and nodes in a tree as either ancestral (blue) or derived (orange) given a labeling of the leaf nodes. An internal node is a derived coalescence if and only if all its descendant leaves are derived. The parent node of the oldest derived coalescence is the mixed coalescence node (represented in black). All other coalescence events are ancestral coalescences. A branch represents a derived lineage if and only if it has a derived node as an ancestor. The mixed lineage is the immediate parent branch of the oldest derived coalescence (black dashed line). All other branches represent ancestral lineages.

Given our labeling, we begin at the present and move back in time, keeping track of the number of derived lineages *n*_*D*_ and ancestral lineages *n*_*A*_ that are present at the given time point. At time points younger than the mixed coalescence node, lineages can only coalesce with other lineages of the same allelic class. The instantaneous coalescence rate within the derived class is *X*_*t*_*N*, and the instantaneous coalescence rate within the ancestral class is s (1 − *X*_*t*_)*N*, where *X*_*t*_ is the frequency of the derived allele at time *t*. At time points older than the age of the mixed lineage, after which there can be no derived lineages, we only consider coalescence within the ancestral class. A similar structured coalescent approach is used by the simulation softwares `discoal` (Kern and Schrider, 2016) and `msprime` 1.0. (Baumdicker et al., 2021)

To formalize the emissions this model emits, given the model enters time *t* with *n*_*D*_ derived lineages and *d* derived coalescences are observed in the interval (*t, t* + 1) at times *t*_1_ to *t*_*d*_, the derived coalescence emission for frequency bin *k* is computed as:

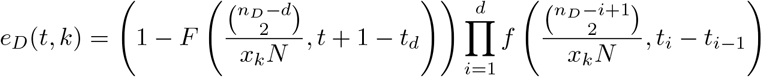

where *f* (*λ, x*) is the density function of an *Exp*(*λ*) random variable at the point *x* and *F* (*λ, x*) is the corresponding CDF. This is the probability of observing the given coalescences (possibly none) at the observed times and then not observing another coalescence before the end of the interval (*t, t* + 1). We compute the ancestral coalescence emissions *e*_*A*_(*t, k*) in the same way, replacing *x*_*k*_*N* with (1 − *x*_*k*_)*N* and considering the number of ancestral lineages *n*_*A*_ and the number of ancestral coalescences *a* in the interval (*t, t* + 1), meaning in total our emission for this timestep is *e*(*t, k*) = *e*_*D*_(*t, k*)*e*_*A*_(*t, k*).

The exceptions to the above expression are the following. If *n*_*A*_ = 1 and *n*_*D*_ = 0, *e*(*t, k*) = 1 if *k* = 1 and *e*(*t, k*) = 0 otherwise. If *n*_*A*_ *>* 1 and *n*_*D*_ = 0, then *e*_*A*_(*t, k*) is calculated as usual if *k* = 1 and *e*_*A*_(*t, k*) = 0 otherwise. If *n*_*A*_ = *n*_*D*_ = 1, then an ancestral emission with *n*_*A*_ = 2 is emitted if *k* = 1 (signalling that the derived allele is no longer segregating) and *e*(*t, k*) = 1 otherwise. Similarly, if *n*_*A*_ *>* 1 and *n*_*D*_ = 1, then an ancestral emission is emitted as usual if *k* ≠ 1 and an ancestral emission with *n*_*A*_ + 1 is emitted if *k* = 1. We find that this framework models the mutation origin and the mixed lineage properly (Figure 3a). We note that the exact handling of the mixed lineage and the formula for the emissions produced by coalescences differs from that described in the original CLUES method.

**Fig 3:**
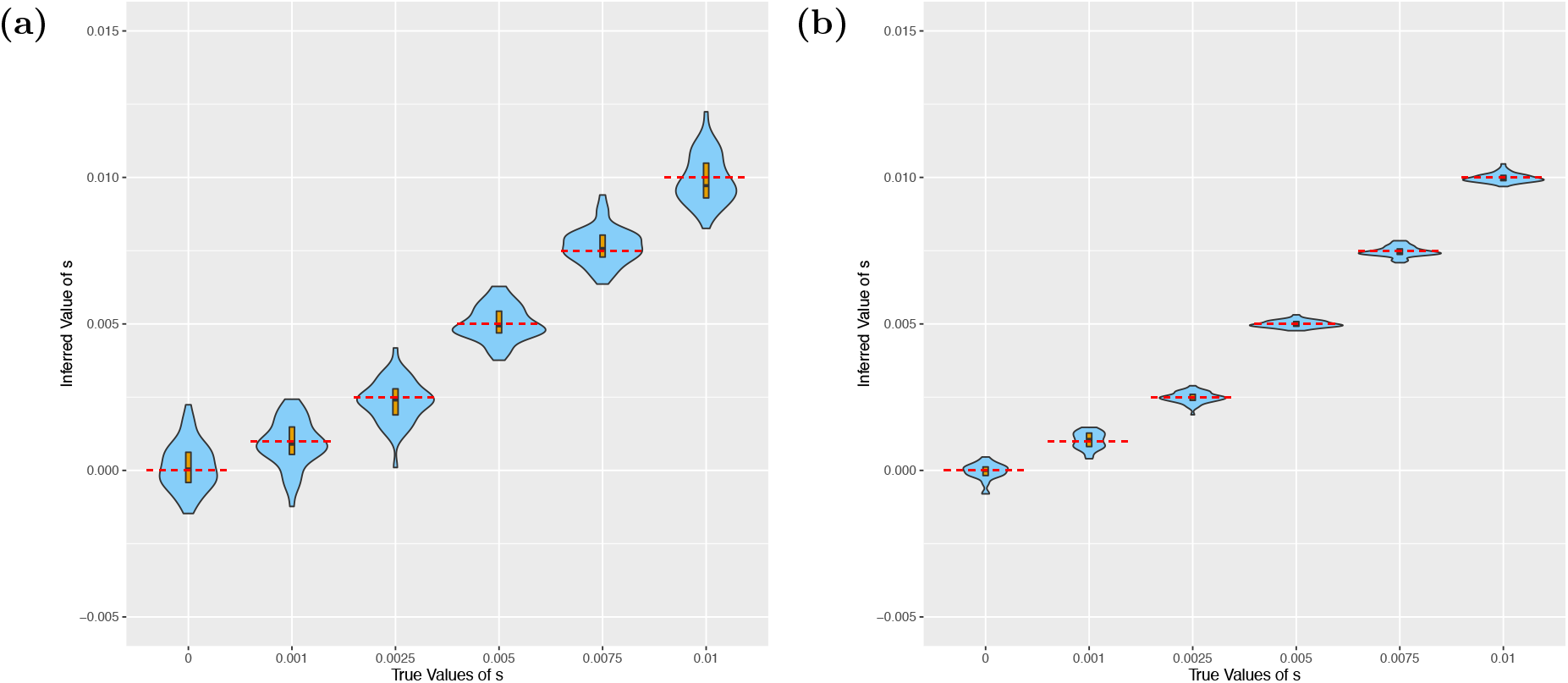
Violin plots showing the results of running CLUES2 on true trees. Boxplots are overlayed with the whiskers omitted. True values of *s* are shown as dashed red lines. 30 replicates were performed for each true value of *s*. Simulations were run with (a) *N* = 30000 and 80 sampled leaves and (b) *N* = 600000 and 800 samples leaves.

There are again two sources of statistical noise in this estimation, which both contribute to the variance

1. Fluctuations in the true allele frequency trajectory due to genetic drift, which causes the true allele frequency to be less informative of the true value of *s*
2. Noisy estimation of the true allele history due to the finite number of lineages sampled.

Therefore, we ran simulations with very low drift and a very high number of sampled leaves (Figure 3b). We observed that the estimated values of *ŝ*^*MLE*^ converged to the true values, which indicates the correctness of our implementation and that the model assumptions we make (such as discretizing time and allele frequency) are sufficiently accurate to not cause substantial biases.

### Importance Sampling of Trees

Of course, the true topology is never known with 100% certainty for real sequence data. Rather, the observed data consists of a set of SNPs around a locus of interest, meaning that the ancestral recombination graph (ARG) must be estimated and one must integrate over samples of the tree at the locus of interest in order to obtain an estimate of the likelihood function. In particular, (Stern et al., 2019) showed that under the assumption that the ARG *G* is independent of *s* given the marginal tree *G*_*k*_ at the SNP of interest, the likelihood ratio 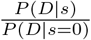 is equal to

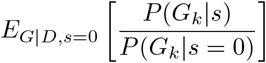

By sampling *M* graphs *G*^1^, …, *G*^*M*^ from the distribution of ancestral recombination graphs given *D* and given *s* = 0 and extracting the marginal tree at our SNP of interest from each graph, one can obtain the following Monte Carlo estimator of the likelihood ratio

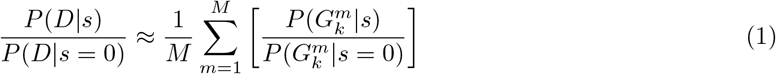

which can be computed by sampling many trees and, for each sampled tree, computing the probability of that tree using the expression derived in the previous section. This reweighting of variates sampled from a different distribution is a technique known as *importance sampling*. It is important to note that this approach adds two additional sources of error, in addition to those described previously.

1. Lack of convergence of the Monte Carlo estimator due to finite *M*.
2. Error incurred by improper sampling of ARGs from the specified posterior distribution. This is to say that the sampled graphs *G*^1^, …, *G*^*M*^ are not *iid* samples from the distribution *P* (*G*|*D, s* = 0). This can happen due to practical considerations, such as poor mixing of the MCMC method used to generate these samples, or poor calibration of the ARG-inference methods themselves.

In particular, with regards to the second possible source of error, we highlight the work of (Brandt et al., 2022) in showing the degree of miscalibration of different ARG-inference methods and how this depends on parameters such as recombination and mutation rate. Therefore, we develop our own MCMC algorithm for sampling gene trees on sequence data in the absence of recombination (see the “Importance Sampling” section of the “Simulation Details” Appendix for more information). This enables us to validate the correctness of our importance sampling implementation without being affected by possible biases induced by improper sampling of ARG-inference methods. We demonstrate the behavior of our importance sampling estimator through a set of illustrative simulations. We begin by simulating genetic data in `msprime` on a set of 24 haplotypes for a 1Mb region with no recombination and with a mutation rate of 7 × 10^−7^. We then run our purpose-built MCMC sampler to generate samples of gene trees given the observed genetic data. We take 1 sample of a tree (M=1) and use it as input to CLUES2. This corresponds to not using the importance sampling framework at all but instead regarding this 1 sampled tree as the true gene tree at this locus with no uncertainty. We find that the mutation rate is so high that it overwhelms the prior centered on *s* = 0 and this 1 sample of a tree does indeed behave like the true gene tree for this data in terms of accuracy (Figure 4a). Then, we run simulations under identical parameter settings (including using *M* = 1 with no importance sampling) except that a mutation rate of *μ* = 2 × 10^−9^ is used. In this case, the estimation of *s* is biased as there is not enough data to overwhelm the prior centered at *s* = 0. This effect is particularly pronounced for larger true values of *s*, which are the values that differ the most from the prior (Figure 4b). This bias can be considered an extreme case of the estimation error that can happen due to small *M*, and it is important to note that this results in bias, rather than increased variance. We note that this bias towards *s* = 0 for small values of *M*, in addition to poor calibration of existing ARG-inference methods, could explain previous analyses that showed a bias of CLUES towards *s* = 0 (see Fig. 4 of (Hejase et al., 2021), Fig. 2 of (Temple et al., 2023)). We also note that using a value of *K*, the number of allele frequency bins, that is too small can result in bias towards *s* = 0.

**Fig 4:**
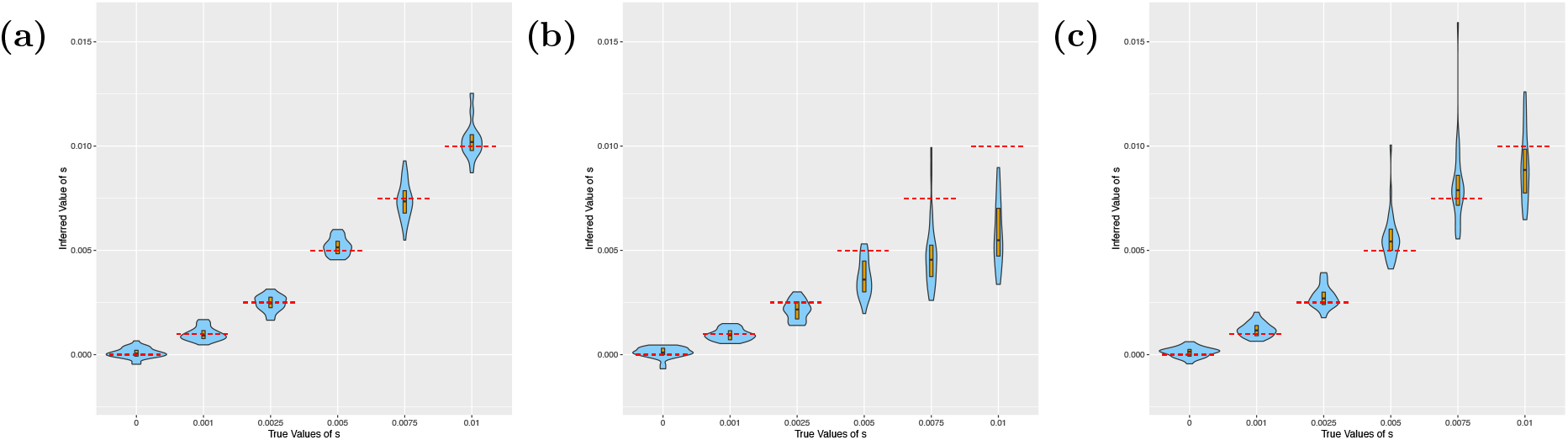
Violin plots showing the results of running CLUES2 on inferred topologies. Boxplots are overlayed with the whiskers omitted. True values of *s* are shown as dashed red lines. 30 replicates were performed for each true value of *s*. Simulations were run with (a) *μ* = 7 *×* 10^−7^ and 1 sample taken without importance sampling, (b) *μ* = 2 *×* 10^−9^ and 1 sample taken without importance sampling, and (c) *μ* = 2 *×* 10^−9^ and 3000 samples taken and used in the importance sampling framework.

To show the utility of our importance sampling approach, we run the same simulation as before with *μ* = 2 × 10^−9^ but instead take *M* = 3000 samples to be used in CLUES2. We find that by properly reweighting the samples of gene trees through our importance sampling framework, we recover approximately unbiased estimates of the selection coefficients (Figure 4c), which validates the implementation of our importance sampling approach.

### Ancient ARGs

Recently, there has been significant interest in using the increasing number of high-quality ancient genomes to infer historic patterns of selection. However, while the usage of time series data has the potential to greatly improve estimates of selection, most existing methods only analyze historic genotype data and do not incorporate information from linked SNPs. (Malaspinas et al., 2012; Mathieson and Terhorst, 2022; Le et al., 2022) In order to fully utilize all the information present in available ancient data, however, it would be desirable to have a method that can incorporate linkage information from SNPs around a locus of interest. To this end, we implement the usage of ancestral recombination graphs on ancient genomes in CLUES2 to incorporate both temporal data and linkage data around the focal site. The general HMM framework remains the same, but the possible emissions of the algorithm change. The input data now consists not only of a list of derived coalescence times and ancestral coalescence times, but also a list of sampling times of derived leaves and a list of sampling times of ancestral leaves. If, for a given time *t*, there are no leaves sampled in the interval (*t, t*+1), then our emissions are the same as for the case of modern ARGs, *e*(*t, k*) = *e*_*D*_(*t, k*)*e*_*A*_(*t, k*). However, if we sample *m*_*D*_ derived leaves in this interval and *m*_*A*_ ancestral leaves in this interval, then our emission becomes 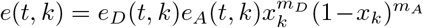 to account for the genotype observations of the leaves. Note that we assume that genotypes are hard-called and do not allow the usage of genotype likelihoods for the leaves of the ARG. After each timestep, we also increase the current number of derived lineages and ancestral lineages as necessary, corresponding to any sampled leaves. To illustrate the correctness of this approach, we simulate ARGs on ancient data under different selection coefficients (see the supplementary section “Simulation Details” for our simulation methodology) and find that we obtain approximately unbiased estimates of the selection coefficient (Figure 5a). Note that while these specific simulations use the true gene trees, importance sampling on ancient ARGs is also possible. To show the improvements in accuracy obtained by utilizing this approach, we also ran CLUES2 using only the modern ARGs obtained from these data and utilizing the ancient data only as genotype emissions, rather than incorporating them into the ARG (Figure 5b). We see that while the estimates of selection are still approximately unbiased, the variance in the estimation is larger due to the smaller amount of linkage information being used in the analysis. Therefore, we highlight the ability of this approach to utilize both time series and linkage information to generate the most accurate possible estimates of selection coefficients.

**Fig 5:**
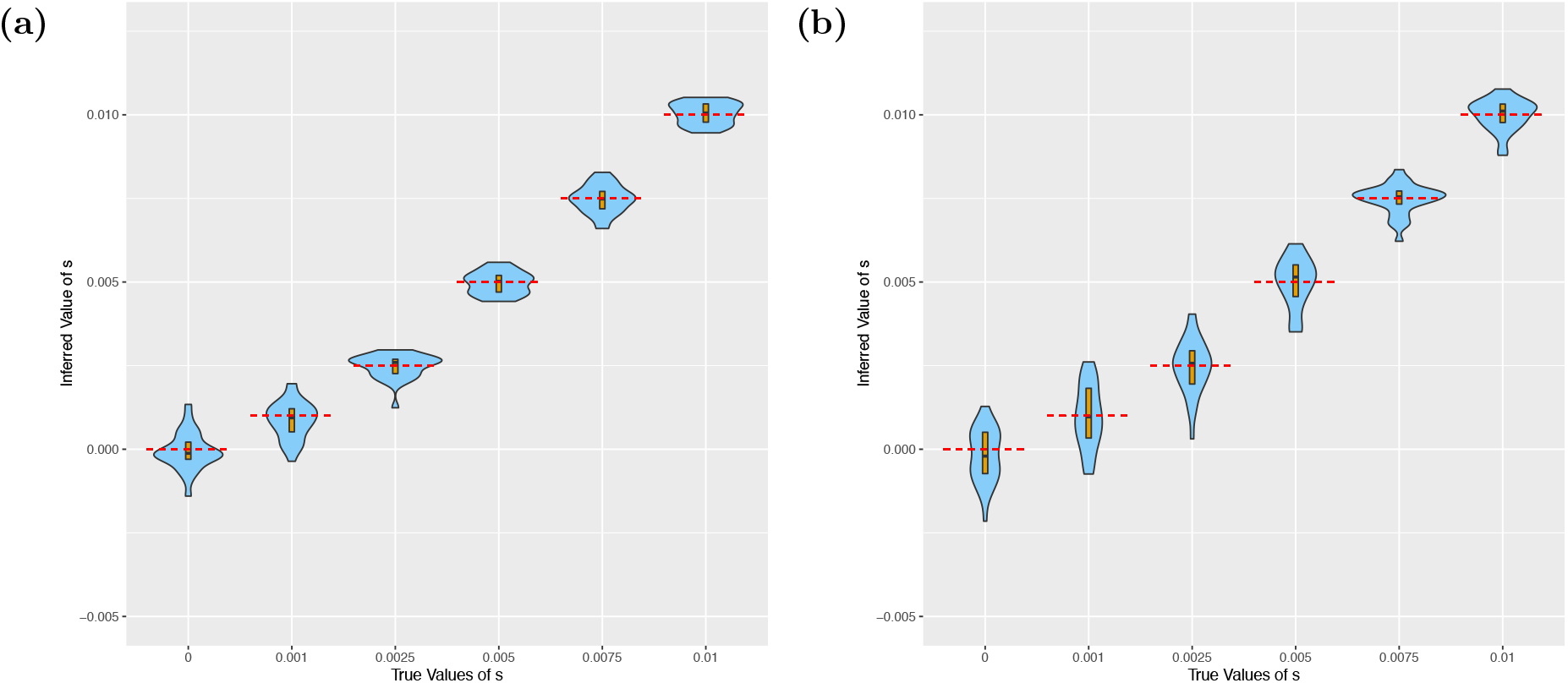
Violin plots showing the results of running CLUES2 on a combination of modern and ancient data. Boxplots are overlayed with the whiskers omitted. True values of *s* are shown as dashed red lines. 30 replicates were performed, with each replicate generating an estimate for each of the 3 selection coefficients. We run simulations (a) where the ancient data is incorporated into the tree and (b) where the ancient data is treated only as genotype emissions.

### Selection in multiple epochs

We also provide functionality for the joint inference of selection coefficients that differ between time periods. This is done by changing the mean of the normal distribution for the transition probabilities from *x*_*k*_ − *sx*_*k*_(1 − *x*_*k*_) to *x*_*k*_ − *s*_*t*_*x*_*k*_(1 − *x*_*k*_) where *s*_*t*_ depends on the timestep *t*. In particular we allow time breakpoints *τ*_1_, …, *τ*_*n*_ to be specified, which results in *n* + 1 different selection coefficients being fit, one for each of the epochs [0, *τ*_1_), [*τ*_1_, *τ*_2_), …, [*τ*_*n*_, *T*). To validate our approach, we simulated ancient genotype data from a model where the selection coefficient in the epoch [0, 200) was 0.01, the selection coefficient in the epoch [200, 600) was 0 and the selection coefficient in the epoch [600, inf) was −0.005. Population size is held constant at 2*N* = 300000, meaning that there is very little drift in the population. We sample 1 diploid individual every generation, and their genotype is recorded. We find that we are accurately able to jointly infer these 3 different selection coefficients (Figure 6).

**Fig 6:**
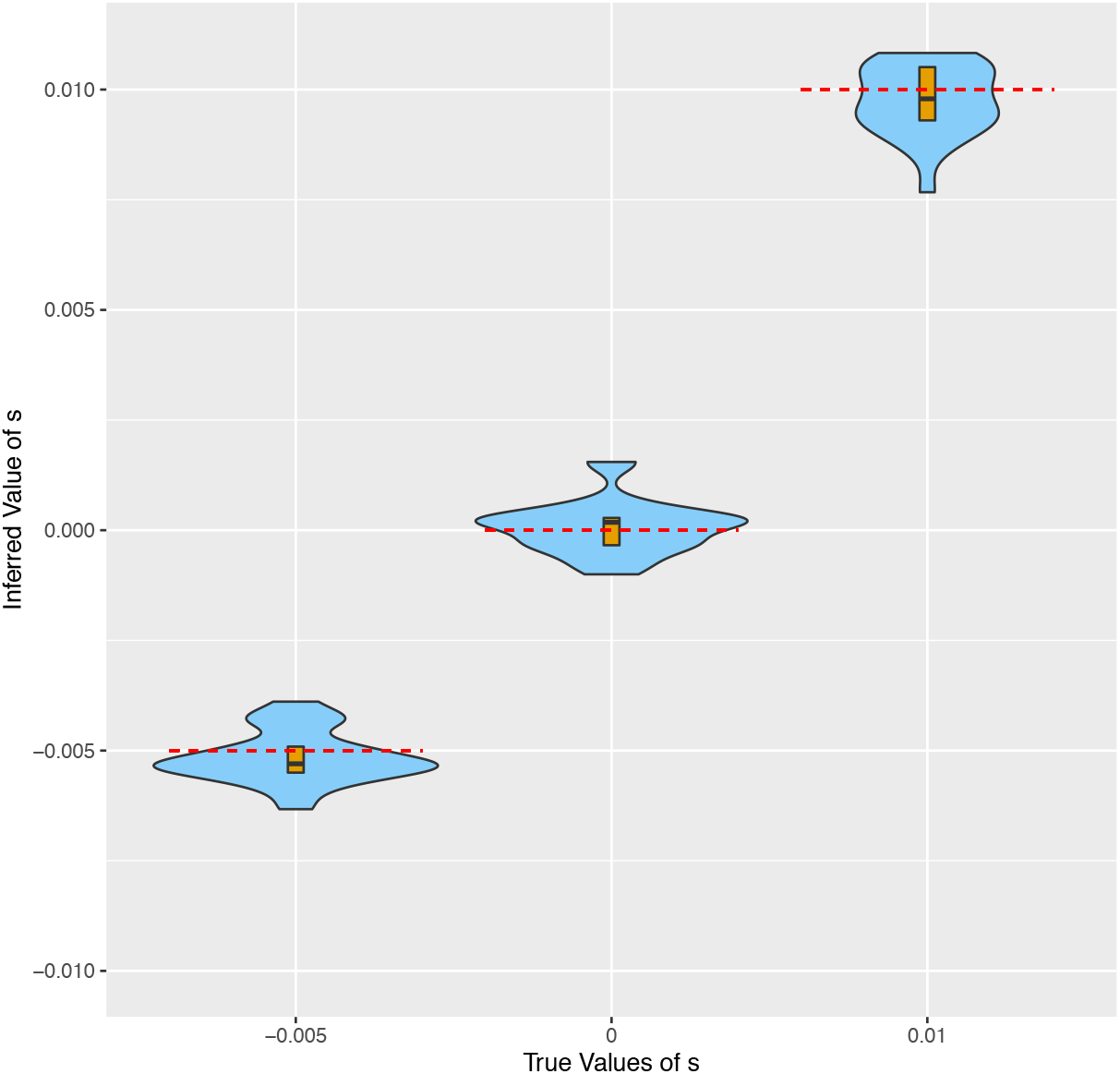
Violin plots showing the results of running CLUES2 on genotype data simulated with differing selection coefficients through time. Boxplots are overlayed with the whiskers omitted. True values of *s* are shown as dashed red lines. 30 replicates were performed, with each replicate generating an estimate for each of the 3 selection coefficients.

### Reconstructing Historic Allele Frequencies

In addition to the inference of selection coefficients and log-likelihood ratios of selection, CLUES2 also has the capability to reconstruct historic allele frequency trajectories by calculating, for each time *t*, the posterior distribution over allele frequencies given the estimate of *s* and the data *D*. This is accomplished by running the backward algorithm, the complement to the forward algorithm, which consists of calculating a matrix *B* following the recursion

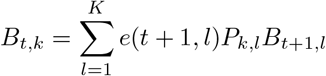

where the last column is initialized to all ones. Then, the desired posterior on allele frequency for a given time point *t*, call it *π*_*t*_, has the form *π*_*t*_(*k*) ∝ *F*_*t,k*_*B*_*t,k*_.

In the original CLUES algorithm, the full forward-backward algorithm is run with the only value of *s* considered being the estimate of *ŝ*^*MLE*^. However, we find that this does not incorporate the uncertainty inherent to the estimation of *s* and therefore results in posterior distributions that are underdispersed. We instead wish to compute the posterior distribution on allele frequencies *P* (*X*_*t*_ = *k*|*D*) assuming a uniform prior on *s*. With a uniform prior on *s*, this distribution is:

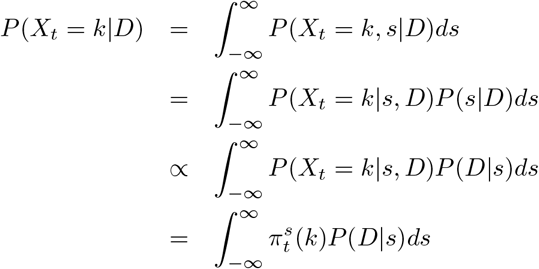

In theory, one could sample a set of discrete values for *s*, call them *s*_1_, …, *s*_*N*_, and approximate this integral using Simpson’s rule, numerical quadrature, or any other way for deterministically approximating the area under the curve of interest. However, these methods scale poorly to higher-dimensional spaces, which will be the case when we are fitting multiple selection coefficients. For this reason, we choose to use Monte Carlo integration, as the variance in the estimate of the integral does not depend on the dimension of the state space but is instead always proportional to 1*/M* where *M* denotes the number of points used in the estimation, (see e.g. (Jarosz, 2008)). Our Monte Carlo estimator is

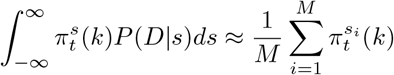

where the *s*_*i*_ are sampled from *P* (*D*|*s*). To do this, we assume that *P* (*D*|*s*) is approximately normally distributed with mean equal to *ŝ*^*MLE*^ as computed by our optimization algorithm. This approximation is based on the Bernstein-von Mises theorem, which states that asymptotically, the posterior is normally distributed with a variance given by the Fisher information matrix. Therefore, in the limit of large data and many Monte Carlo samples, this approximation becomes exact.

Using the pairs of selection coefficients and values of the likelihood function *P* (*D*|*s*) computed during our optimization routine, call them (*s*_*j*_, *L*(*s*_*j*_)), we then fit the variance (or covariance matrix) to our data points using a least-squares regression model. Concretely, we find the value of Σ that minimizes

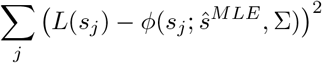

where *ϕ*(*x*; *μ*, Σ) is the density of a normal distribution with mean *μ* and covariance matrix Σ evaluated at the point *x*. We then sample a number of points *M* from this distribution (10*d* by default, where *d* is the dimension of *s*) and compute 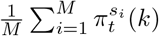 as our estimate of the posterior distribution over allele frequencies at time *t*. In this way, we take uncertainty in our estimation of *s* into account for our estimate of the posterior distribution of allele frequencies. We validate this approach by comparing it to an exact rejection sampling approach for reconstructing historic allele frequencies. Specifically, we fix a set of genotype observations and run CLUES2 with our Monte Carlo integration approach. Then, we repeatedly sample selection coefficients uniformly from [−0.1, 0.1] and simulate allele histories conditional on our sampled coefficient. We then sample genotype observations conditional on these trajectories and retain only those trajectories associated with datasets that match our fixed set of genotype observations. This is done until we have retained 1000 trajectories, from which we can reconstruct empirical posterior frequency intervals. We compare these two approaches in Figure S5. We also perform a comparison where 2 selection coefficients are independently inferred, where the coefficient in each interval is sampled uniformly from [−0.1, 0.1] (Figure S6). For both simulations, we find strong concordance between our Monte Carlo integration framework and the exact rejection sampling approach.

### Validation of chi-squared test

Along with reporting estimates of selection coefficients, we report the log-likelihood ratio ln 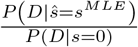 and the associated p-value for selection. By Wilks’ Theorem, if data is simulated under the null hypothesis of *s* = 0, 2 ln 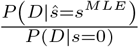 should asymptotically follow a *χ* distribution with degrees of freedom equal to the difference in the dimensions of the parameter spaces of *ŝ*^*MLE*^ and *s* = 0 (Wilks, 1938). As *s* = 0 has dimension 0, the degrees of freedom is always equal to the number of different selection coefficients that are estimated in different epochs (which is 1 by default). Furthermore, the p-value obtained from this chi-squared distribution should be uniformly distributed on [0, 1] under the null hypothesis.

We therefore perform validations on each of our above simulations by examining the distribution of 2 log(*LR*) when data is simulated under *s* = 0 to the expected chi-squared distribution with *k* degrees of freedom where *k* is the number of independent selection coefficients that are estimated (3 in the section “Selection in multiple epochs”, 1 in all other simulations). We compare the quantiles of the empirical distribution of 2 log(*LR*) to the theoretical chi-squared distribution using a P-P plot. We also plot a histogram with 5 bins of the p-values obtained from these log-likelihood ratios by using the upper tail of the appropriate chi-squared distribution and comparing it to the uniform distribution. The results are shown in Supplementary Figures S7 through S12. We find that for each simulation we consider, the empirical distribution of 2 log(*LR*) follows the expected chi-squared distribution and that the corresponding p-values are distributed uniformly between 0 and 1, thus indicating that CLUES2 is properly calibrated and can be used in rigorous hypothesis testing frameworks.

### Computational Improvements

We make modifications to the naive forward-backward algorithm that significantly improve the computational runtime of CLUES2. In practice, there are two steps of the basic forward algorithm implementation of CLUES2 that require significant computational runtime. The first is the computation of the *K* × *K* per-generation allele frequency transition matrix *P*. Naively, this requires the computation of *K*^2^ entries. However, many of these probabilities will be close to 0 as the per-generation probability of transitioning from a high allele frequency to a low allele frequency in one generation and vice-versa is very small. More formally, we observe that the nature of the Gaussian transition probabilities ensures that the transition matrix is a sparse banded matrix where appreciable probability only falls near the main diagonal. Therefore, we instead, for each allele frequency bin *k*, compute only the entries in the matrix corresponding to the probability of transitioning to frequencies lying in the range [max(0, *k* − *ϵ*), min(1, *k* + *ϵ*)]. In practice, we set *ϵ* = 0.05. For frequencies from *ϵ* to 1 − *ϵ*, this means that the possible frequencies in the next generation fall in a range representing 10% of the frequency space, meaning that the computational speedup should be approximately 10x for this step. We note that this idea of approximating the full per-generation transition matrix by a sparse banded matrix has been previously applied to the full Wright-Fisher model (Spence et al., 2023). We call this Approximation A.

The other section that requires significant time is the execution of the forward algorithm, which must be run every time the likelihood function is called during the optimization procedure (or *M* times for each call to the likelihood function if importance sampling is used with *M* samples). Each call to the algorithm integrates over all derived allele trajectories from time 0 to time *T*, though the dynamic programming nature of the forward algorithm means that this integration is done in polynomial time. However, some of these trajectories have vanishingly small probabilities, such as the oscillating trajectory where *x*_*i*_ = 0 for *i* odd and *x*_*i*_ = 1 for *i* even. We instead modify the forward algorithm to only integrate over likely allele trajectories. When looking at the forward algorithm equation:

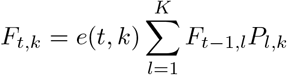

We notice that many of these transition probabilities *P*_*l,k*_ will be close to 0 if *l* is quite far from *k* (and if |*x*_*k*_ − *x*_*l*_| *>* 0.05, the probability will be exactly 0 based on our revised computation of the transition matrix *P*). This means that these values of *l* can be left out of the summation without significantly affecting the sum. Therefore, we compute, for all *K* columns of *P*, a lower row index *a*_*k*_ and upper row index *b*_*k*_ such that the sum of the entries between *a*_*k*_ and *b*_*k*_ is at least 99.9% of the total sum of the entries in column *k*. We then replace the above sum by

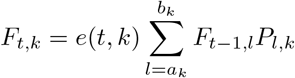

which saves considerable computational time while still retaining accuracy. These lower and upper bounds can be computed at the same time as the transition matrix *P* is computed and do not need to be recomputed at each timestep of the forward algorithm. Conceptually, this is equivalent to replacing the Gaussian diffusion approximation with a Gaussian distribution that is truncated at extreme values. We call this Approximation B.

We also make another computational improvement based on the observation that when computing

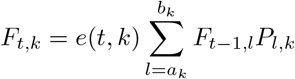

even for indices, *l*, with fairly large values of *P*_*l,k*_, the computed sum might still be near 0 if the values of *F*_*t*−1,*l*_ are quite small. Therefore, we choose to simply not compute values of *F*_*t,k*_ if we can reasonably conclude that they will be near 0. We do this by calculating a lower bound *α*_*t*_ and an upper *β*_*t*_ between which most of the probability of the column *F*_*t,·*_ lies (i.e. bounds for which 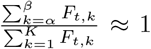). We then compute the entries *F*_*t*+1_, *k* only for the *k* satisfying *α*_*t*_ − *K/*20 *< k < β*_*t*_ + *K/*20, where the *K/*20 terms are chosen to correspond to the *ϵ* = 0.05 restriction on the transition matrix. The other entries in that column of *F* are left as 0. *α*_*t*+1_ and *β*_*t*+1_ are then set respectively to the minimum and maximum indices *k* for which 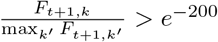. We set *α*_1_ and *β*_1_ to both be equal to the index of the row corresponding to the allele frequency bin closest to the observed modern allele frequency. Informally, we are assuming that each column of *F* is relatively close in distribution to the previous column of *F*. This is a reasonable assumption if the emissions at a given timestep do not significantly change the probability distribution across the hidden states. The only emission that does violate this assumption is the coalescence of the mixed lineage, as we enforce the fact that the derived allele frequency must be 0 at this time (as we described in section “True Trees”). For this reason, if we observe only 1 derived lineage left at time *t*, we set *α*_*t*_ to 1, so we always compute *F*_*t*,1_ and all values *F*_*t,k*_ for *k < β*_*k*_ and are therefore never “surprised” by observing this coalescence. We find that this approximation saves significant computational time and has a negligible effect on the estimated value of the selection coefficient. This practice of running the forward algorithm while only keeping track of a smaller number of “best” states is inspired by the beam search approaches used to approximate the Viterbi decoding of an HMM (Deshmukh et al., 1999), although the exact details differ. We call this Approximation C. The concepts underyling Approximations A, B, and C are shown in Figure 7.

**Fig 7:**
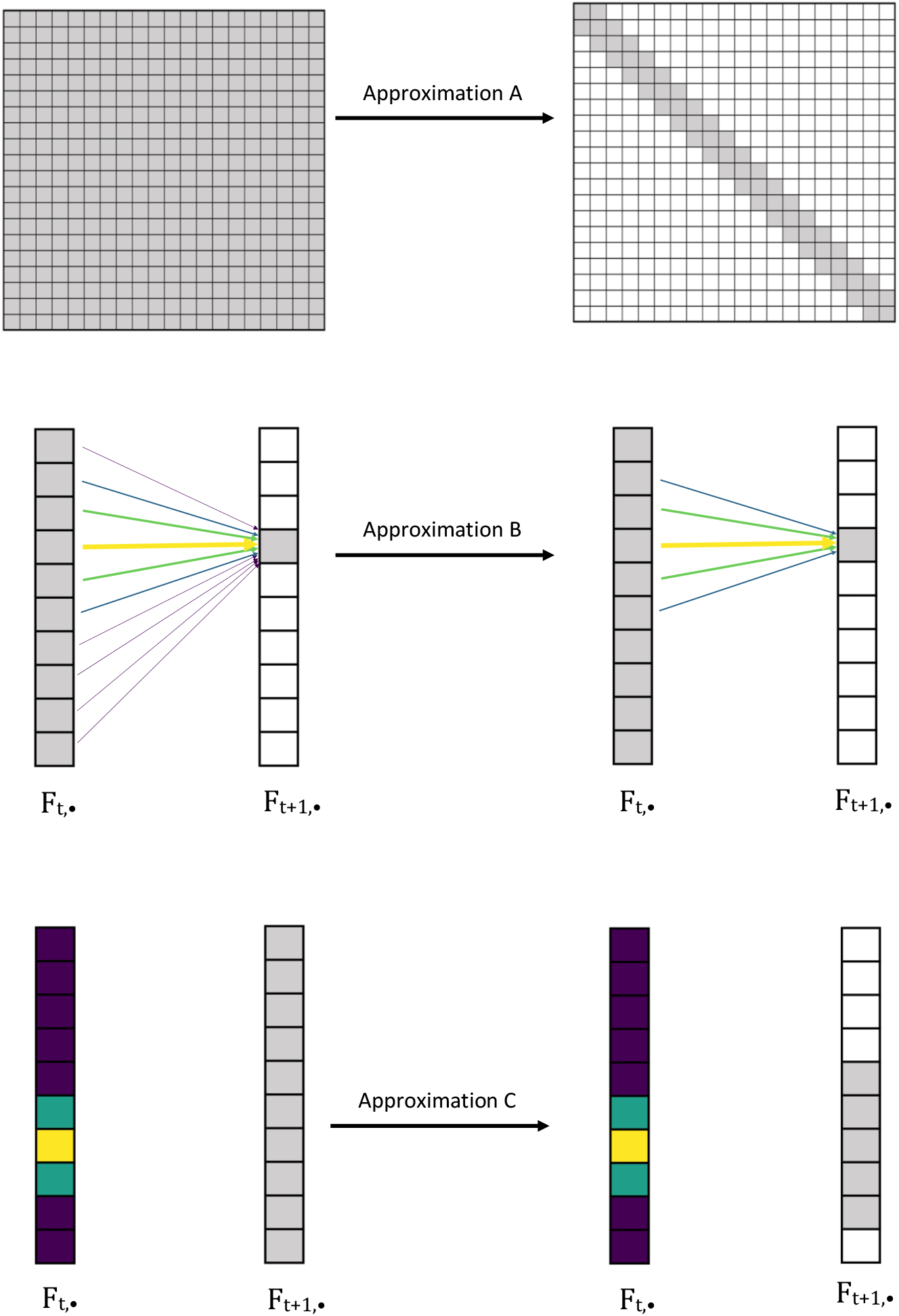
Illustration of Approximations A, B, and C. Approximation A approximates the transition matrix by a sparse banded matrix. Approximation B reduces the number of states in the previous column of *F* that are summed over to compute each entry of the forward matrix *F*. Approximation C reduces the number of entries that are computed in a column of the forward matrix *F* based on the probability density of the previous column of *F*. Here, the gray entries represent values that are computed, while the colored entries and arrows represent transition or forward probabilities. Lighter colors denote higher probabilities, and darker colors denote smaller probabilities.

For the backward algorithm, we also perform Approximation A, and make an analogous approximation to Approximation B, where *a*_*k*_ and *b*_*k*_ instead represent indices for which the sum of the entries between *a*_*k*_ and *b*_*k*_ is at least 0.999 of the total sum of the entries in *row k*, as opposed to column *k*. We do not make an analogous approximation for Approximation C. In practice, this set of computational improvements greatly improves the runtime of CLUES2 compared with the original CLUES algorithm. We show this by measuring the runtime of both CLUES and CLUES2 as a function of the number of importance samples used for a given dataset. We plot the results in Figure 8.

**Fig 8:**
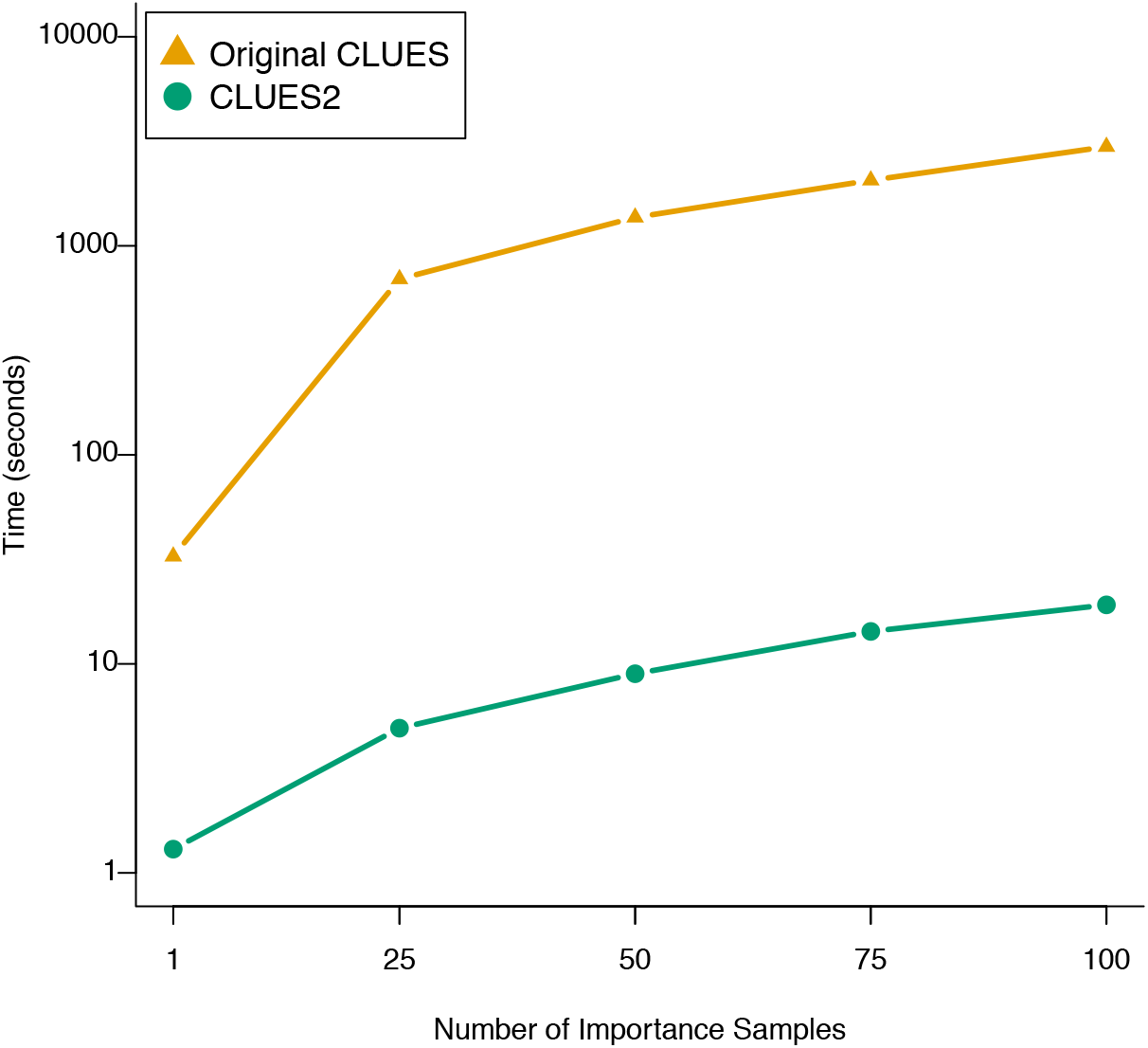
Comparative runtime of CLUES and CLUES2 on different numbers of importance samples.

We see that CLUES2 is over 10 times faster when considering only 1 importance sample and is over 100 times faster when 100 importance samples are used. These computational speedups are important for several reasons. Firstly, they allow more importance samples to be used for the same amount of computational resources, which contributes to better convergence of the Monte Carlo estimator of the log-likelihood ratio (Equation 1) and thus improved accuracy. Secondly, it expands the applicability of CLUES2 to large numbers of SNPs, for example in genome-wide scans of selection. We show that these approximations have a negligible impact on our estimate of the log-likelihood function by comparing results both with and without these approximations (see Figure 9).

**Fig 9:**
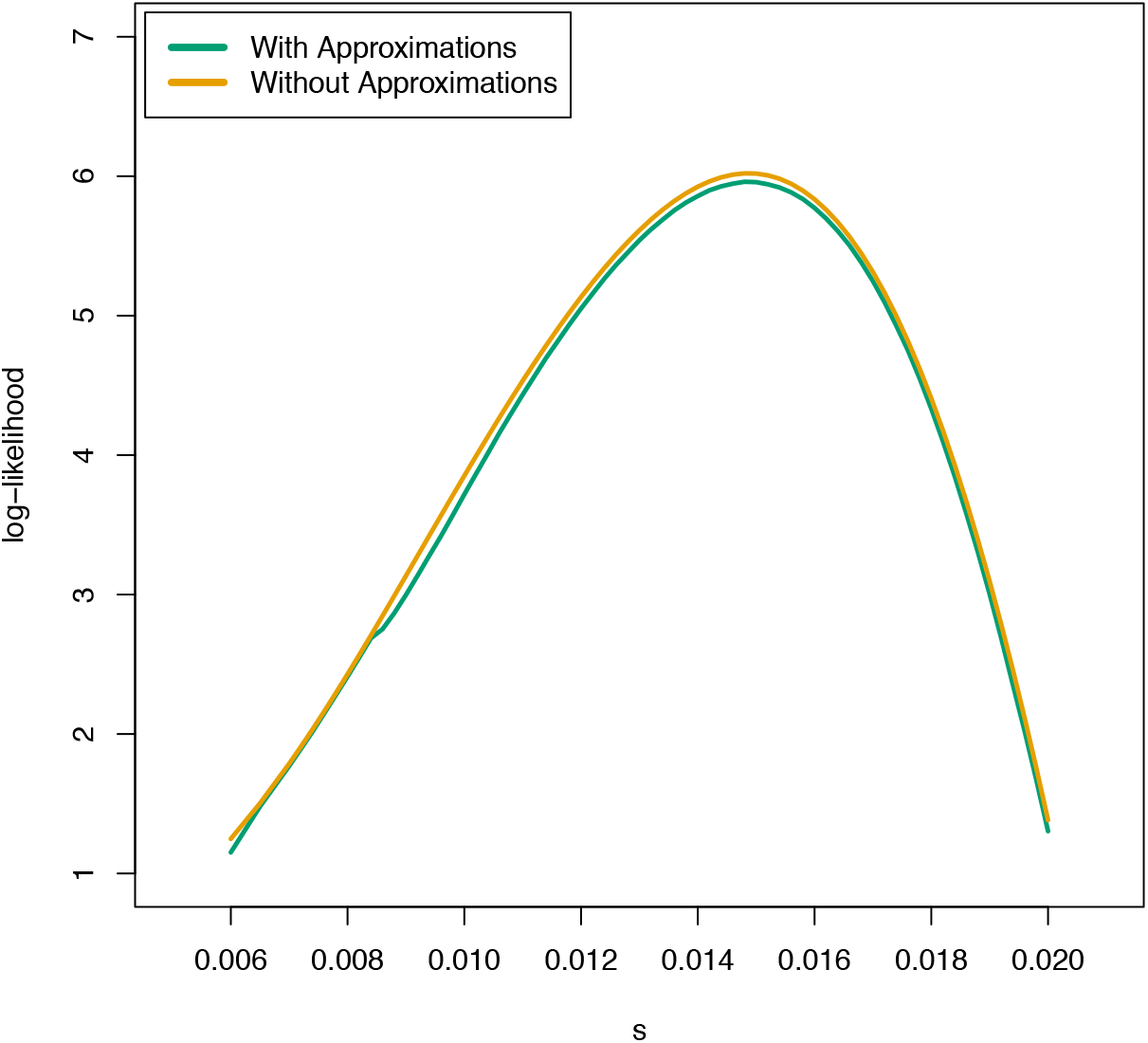
Comparison of the estimates of the log-likelihood function of our dataset of 100 importance samples both with and without our HMM approximations. The log-likelihood was evaluated at 60 values of *s* spaced equally between 0.006 and 0.02 for each case, and the plots of the functions were generated via linear interpolation.

### Inferring Gene Trees on Ancient Human Data

We applied CLUES2 to a recently published collection of imputed and phased ancient genomes (Allentoft et al., In Press). This dataset consists of 1664 diploid individuals sampled from around the world and has both newly sequenced genomes as well as previously published genomes. The highest concentration of these genomes is located in Western Eurasia, and for that reason, we restricted our analysis to West Eurasian genomes. In addition, it has been shown that inferring ARGs on very low coverage data can cause biases in the estimation of pairwise coalescent times (see Supplementary Section 2 of (Allentoft et al., In Press)), so we chose to limit our analysis to a subset of 187 sampled West Eurasian genomes with a coverage of at least 2x. We then merged this ancient data with a dataset of 100 randomly chosen individuals from the 1000 Genomes Phase 3 EUR Superpopulation, resulting in 287 total diploid individuals (Consortium, 2015).

To analyze a chosen set of SNPs, we ran Relate (Speidel et al., 2019) on the corresponding chromosomes of our merged dataset, using recombination maps and genomic masks from the 1000 Genomes Project Phase 3 in addition to the GRCh37 human ancestral sequence (Consortium, 2015) to polarize the alleles. The `EstimatePopulationSize` capability was used in order to generate a .coal file representing the estimated historic population sizes, which was then used as input to the main Relate algorithm. The `SampleBranchLengths` function was used to generate 2000 samples of the marginal tree at each focal SNP, and we converted this output to the CLUES2 input using a custom Python script `RelateToCLUES.py`. For certain SNPs, the sampled topologies did not satisfy the infinite sites assumption with respect to the focal SNP. When this occurs, we performed the minimum number of “leaf flips” such that the infinite sites assumption is satisfied. A “leaf flip” consists of switching the allele of a leaf node from derived to ancestral or vice versa. Note that while the minimum number of flips is always well-defined, the exact leaves to be flipped is not necessarily unique. If this is the case, one set of leaves to flip is chosen deterministically based on the ordering of leaf nodes in the given Newick representation. We list the number of leaf flips performed for each SNP in Table 2. We than ran CLUES2 on each of these input files with the arguments `--df 600`, `--tCutoff 536` (which, assuming a generation time of 28 years, (Moorjani et al., 2016) corresponds to 15000 years) and a value of `--popFreq` that was estimated from our 100 modern individuals.

## Results

### Comparison with Existing Methods

A variety of existing methods have been used to identify selection coefficients. It is not directly possible to test all features of CLUES2 against these methods as to our knowledge CLUES2 is the first method that can model changing selection coefficients through time in a hypothesis-testing framework and that can do this on samples of ARGs on ancient data. However, we do compare the results of CLUES2 with other methods on our low-mutation simulated genotype data described in the section “Importance Sampling of Trees” and plotted in Figure 4c. As this data consists of all modern samples and we are only estimating a constant selection coefficient through time, many existing methods are applicable to this data. We consider 3 other methods, all based on summary statistics of the data. The first is Tajima’s D, a site-frequency-spectrum based statistic which is based on the relative difference between the number of segregating sites and the nucleotide diversity from that expected under neutrality. (Tajima, 1989). The second is H12, a haplotype-based statistic that calculates the imbalance in the relative frequencies of the different haplotypes observed in the data. (Garud et al., 2015) The third is nSL, which is based on calculating the average number of segregating sites around a focal SNP that are identical by state among haplotypes with the derived allele, calculating the corresponding average among haplotypes with the ancestral allele, and taking the log-ratio of the 2 quantities. (Ferrer-Admetlla et al., 2014) We measure the ability of these 3 methods to infer selection coefficients through an ABC approach, detailed in the following pseudocode.

#### Algorithm 1

Approximate Bayesian Computation Algorithm

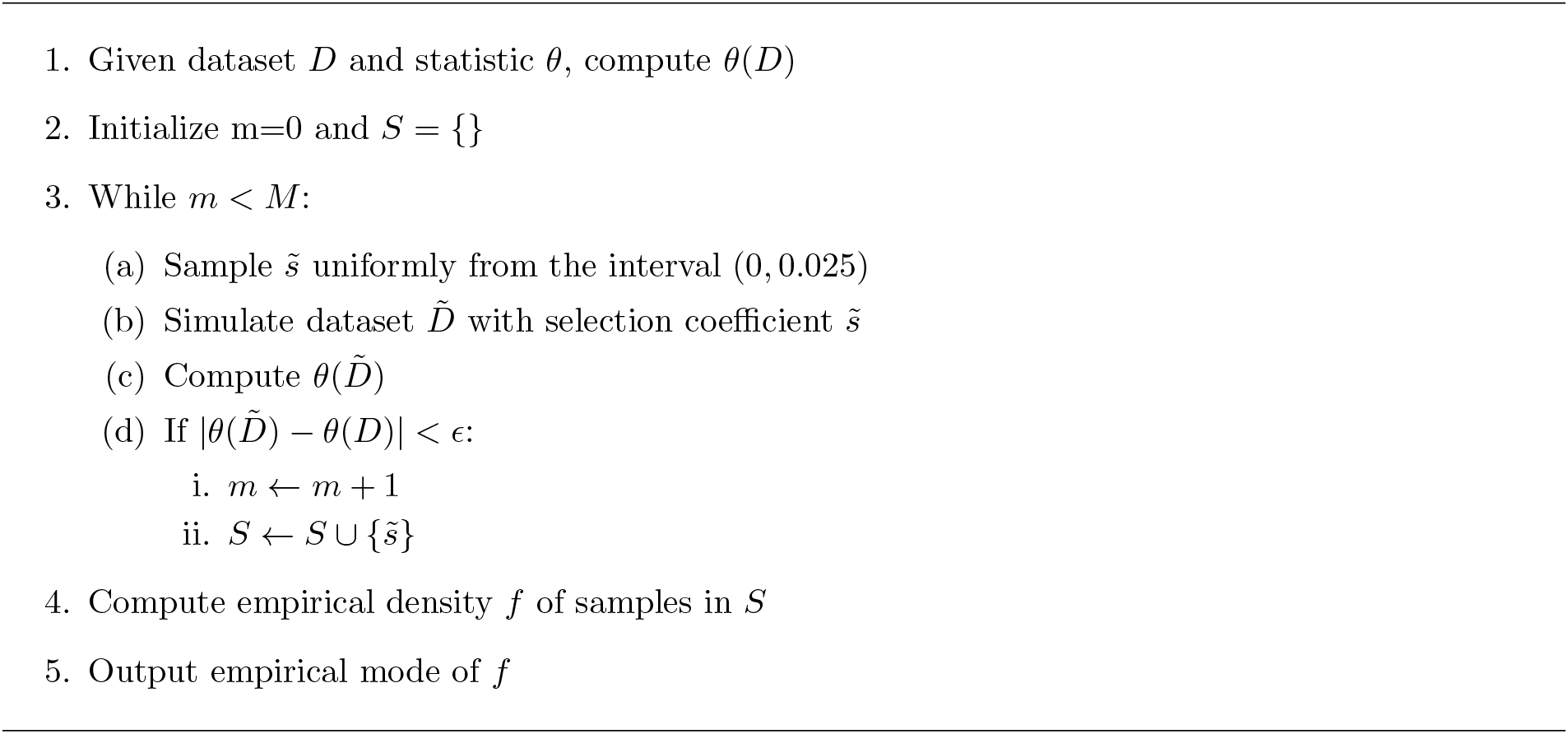

Notice that we use a uniform (0, 0.025) prior, which allows the empirical mode to be interpreted as an approximation to the maximum likelihood estimator. We run this algorithm for each dataset using *M* = 2000 iterations, and *ϵ* values of 0.015, 0.001, and 0.05 for Tajima’s D, H12, and nSL respectively. The posterior is estimated, for each set of resulting ABC sampled values of *s* (*S*), using kernel density estimation with the Epanechnikov (parabolic) kernel and a bandwidth specified by Silverman’s ‘rule of thumb’ (Epanechnikov, 1969; Silverman, 1986). We found that increasing *M* and decreasing *ϵ* past these thresholds did not significantly change the estimates. It is worth noting that because of the relatively small set of values H12 can take for 24 haplotypes (it is bounded above by the partition number p(24)=1575), we can explicitly calculate that the value of *ϵ* = 0.001 reduces this ABC approach to exact rejection sampling. This is to say that two datasets of 24 haplotypes have H12 values that differ by less than 0.001 if and only if their H12 values are exactly the same. We plot the results of this analysis compared with the CLUES2 results in Figure 10. Note that we only analyze the 4 largest selection coefficients as `msprime` cannot simulate sweeps with negative selection coefficients, meaning that edge effects in the density estimation can happen for very small selection coefficients.

**Fig 10:**
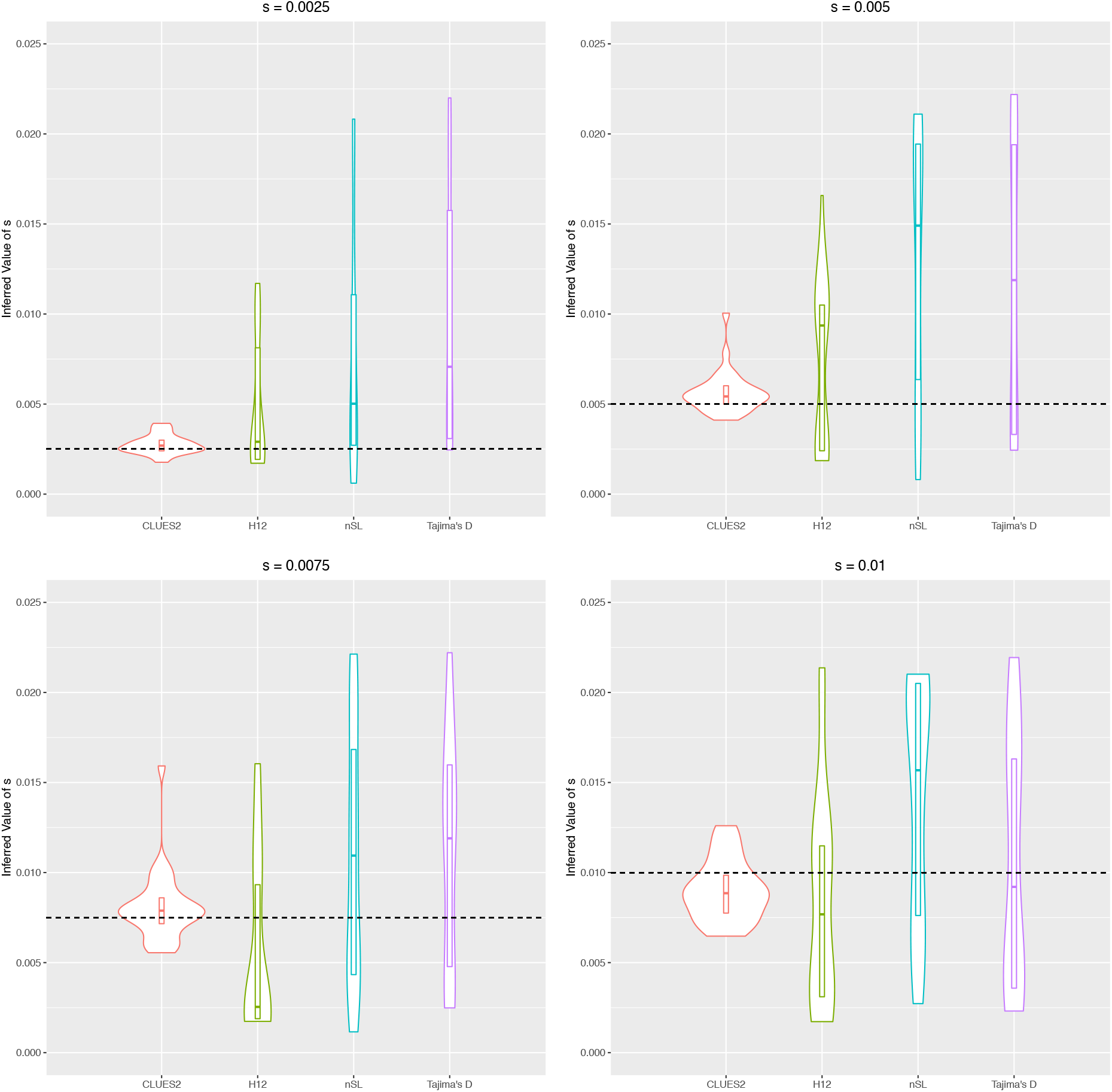
Comparison of the ability of CLUES2 and several summary statistic based methods to infer selection coefficients. We infer selection coefficients from the summary statistics using the ABC algorithm described above. The dataset is identical to that analyzed in Figure 4c. The true simulated values of the selection coefficients are shown as horizontal dashed lines.

We see that all the summary statistic based methods show much greater variance than CLUES2, which is to be expected given the comparative lack of information present in summary statistics when compared with the full dataset.

### Analysis of Ancient Human Data

We chose 4 SNPs to analyze in the ancient human dataset that have previously been identified as candidates for selection in Eurasians. The first is rs4988235, located in the MCM6 gene, where the derived allele is associated with lactase persistence (Enattah et al., 2002; Bersaglieri et al., 2004; Chin et al., 2019). The second is rs35395 in the SLC45A2 gene, where the derived allele is associated with lighter skin pigmentation (Marcheco-Teruel et al., 2014; Tiosano et al., 2016; Lloyd-Jones et al., 2017; Lona-Durazo et al., 2019). The third is rs12153855 in the TNXB gene of the HLA region, where the derived allele is associated with age-related macular degeneration and atopic dermatitis (Cipriani et al., 2012; Weidinger et al., 2013; Grange et al., 2015; Ye et al., 2016). The fourth is rs75393320 in the ACP2 gene, where the derived allele is associated with increased HDL cholesterol (the so-called “good cholesterol”) (Klarin et al., 2018; Ke et al., 2021). General information about each SNP is described in Table 1, where the derived allele frequency is calculated from our 100 sampled EUR individuals from the 1000 Genomes Project.

**Table 1:** General information about the 4 SNPs we considered in our ancient human analysis.

| Gene | SNP rsID | GRCh37 Position | Anc/Der Allele | Derived Allele Freq |
| --- | --- | --- | --- | --- |
| MCM6 | rs4988235 | chr2:136608646 | G/A | 0.535 |
| SLC45A2 | rs35395 | chr5:33948589 | T/C | 0.930 |
| TNXB | rs12153855 | chr6:32074804 | T/C | 0.125 |
| ACP2 | rs75393320 | chr11:47266471 | G/C | 0.150 |

**Table 2:** Results of the 1-epoch model. One selection selection coefficient is estimated that is constant through time. Bolded rows indicate the genes for which this is the best model as determined by having the lowest AIC. We report the number of leaf flips necessary for each SNP. We do not repeat this information in the following tables as it depends only on the inferred tree structure and thus remains consistent across models.

| Gene | $s_{0-15000}^{MLE}$ | $-\log_{10}(p)$ | AIC | Leaf Flips |
| --- | --- | --- | --- | --- |
| MCM6 | 0.00934 | 80.86 | -364.01 | 2 |
| SLC45A2 | 0.00734 | 50.30 | -223.74 | 3 |
| TNXB | 0.01011 | 14.00 | -57.88 | 8 |
| **ACP2** | **0.00461** | **8.14** | **-31.45** | **0** |

**Table 3:** Results of the 2-epoch model. There is one selection coefficient breakpoint at 5000 years before the present. Bolded rows indicate the genes for which this is the best model as determined by having the lowest AIC.

| Gene | $s_{0-5000}^{MLE}$ | $s_{5000-15000}^{MLE}$ | $-\log_{10}(p)$ | AIC |
| --- | --- | --- | --- | --- |
| MCM6 | 0.01244 | 0.00498 | 83.45 | -380.30 |
| SLC45A2 | 0.01434 | 0.00142 | 56.37 | -255.62 |
| **TNXB** | **0.01804** | **-0.00655** | **16.83** | **-73.51** |
| ACP2 | 0.00539 | 0.00367 | 7.32 | -29.73 |

**Table 4:** Results of the 3-epoch model. There are two selection coefficient breakpoints at 2500 and 5000 years before the present. Bolded rows indicate the genes for which this is the best model as determined by having the lowest AIC.

| Gene | $s_{0-2500}^{MLE}$ | $s_{2500-5000}^{MLE}$ | $s_{5000-15000}^{MLE}$ | $-\log_{10}(p)$ | AIC |
| --- | --- | --- | --- | --- | --- |
| **MCM6** | **0.00674** | **0.01988** | **0.00391** | **83.74** | **-385.15** |
| **SLC45A2** | **-0.00214** | **0.03292** | **0.00074** | **58.49** | **-268.55** |
| TNXB | 0.02016 | 0.01441 | -0.00607 | 16.83 | -71.86 |
| ACP2 | 0.00474 | 0.00650 | 0.00349 | 6.65 | -27.757 |

For each SNP, we calculated the maximum likelihood estimate of the selection coefficient, the p-value as computed from the log-likelihood ratio, and the resulting Akaike Information Criterion (AIC) (Akaike, 1974). We ran CLUES2 on each of these SNPs assuming a constant value of the selection coefficients through time, with a selection coefficient breakpoint at 5000 years ago, and with two breakpoints at 2500 years ago and 5000 years ago. We show the results in Tables 2 to 4, where we group the results by the number of selection breakpoints. We also plot our estimates of the derived allele frequency trajectories for each SNP under the model with the lowest AIC in Supplementary Figures S1 to S4.

### Ancestry-stratified Selection Analysis

The selection inference framework of CLUES2 relies on interpreting significant changes in the frequency of an allele as indicative of selection acting on that allele. However, population structure could confound the inference of selection by contributing to the change in the frequency of an allele even in the presence of selective neutrality. In particular, an observed increase in the frequency of an allele in a population could occur in the absence of selection due to either admixture from a source population where that allele is segregating at a higher frequency or due to a correlation between ancestry and sample age. To correct for this, we utilized the local ancestry labelings available for this dataset that classify each SNP in a particular haplotype as belonging to one of 4 major ancestral populations that contribute to the ancestry of modern Europeans: Anatolian farmers (ANA), Caucasus hunter-gatherers (CHG), Western hunter-gatherers (WHG), and Eastern hunter-gatherers (EHG) (Pearson and Durbin, 2023; Irving-Pease et al., In Press). For a given SNP, we then partition each of the ancient haplotypes into these group based on their labelings and compute the modern frequency of the allele in each ancestry group by measuring the percentage of European 1000 Genomes haplotypes of that ancestry which have the derived allele. CLUES2 was then run for each of the 4 ancestries for each SNP using the number of epochs chosen by AIC in our non-stratified analysis described in the previous section. Reconstructed allele frequency trajectories and p-values for the selection tests are plotted in Figures S13 to S16. We highlight that this analysis, in addition to correcting for population structure, only utilizes ancient haplotypes and therefore is not affected by possible biases incurred by the miscalibration of ARG-inference methods.

## Discussion

In this paper, we introduced CLUES2, a flexible, full-likelihood method that is able to utilize more of the information present in sequence data than summary statistics. CLUES2 has the ability to generate unbiased estimates of selection coefficients and also generates well-calibrated p-values in order to run statistical tests for selection. The methodology for our analysis of the West Eurasian dataset highlights the ability of CLUES2 to identify distinct selection coefficients in different epochs. In particular, we emphasize that different hypotheses can be tested for a given SNP without having to obtain new samples of gene trees. Furthermore, the new functionality to analyze ARGs on ancient data enables the usage of both time-series data and information from linked SNPs in selection analyses.

For the ancient human DNA analysis, we observe that the 1-selection coefficient model provides the best fit for the ACP2 SNP in the pan-ancestry analysis, corresponding to a classic selective sweep in progress. As increased levels of HDL have been shown to help reduce the risk of myocardial infarction (heart attack) in contemporary medical patients (Remaley et al., 2008), this result indicates that a similar benefit may have existed in ancient Eurasian populations as well. The ancestry-stratified analysis for this SNP suggests that this sweep may have been localized to WHG ancestry. The 2-selection coefficient model provides a best fit for the pan-ancestry analysis of the TNXB SNP, which we see corresponds to a recent period of positive selection preceded by a period during which the derived allele was slightly disfavored. We speculate that this relatively recent selection on a gene in the HLA region, which is known to be associated with immune function (Gough and Simmonds, 2007), could be the result of increasing population density and exposure to domesticated animals that occurred around this time. (Page et al., 2016; Marciniak et al., 2022) The massive demographic shift induced by the introduction of agriculture likely exposed humans to new selective pressures on genes associated with immune function, explaining why we see selective sweeps on certain immunity-associated alleles beginning during this period. The ancestry-stratified analysis for this SNP suggests that this sweep may have been localized to CHG ancestry. Sparse sampling at very old time points contributes to high uncertainty in the allele frequency of the SNP for many ancestry groups.

We see that the 3-selection coefficient models provide best fits for the pan-ancestry analysis of the other SNPs. For the MCM6 locus, we find a significant increase in the lactase persistence allele beginning approximately 6000 years before the present. We note that this is thousands of years after the consumption of dairy began in Europe, as evidenced by milk fat residues discovered in potsherds dating back to at least 9000 years before the present (Evershed et al., 2022). This gap, which has previously been noted in several studies (Itan et al., 2009; Mathieson et al., 2015; Burger et al., 2020), suggests that the selective pressure for lactase persistence was not in fact the initial domestication of animals and ensuing increase in lactose consumption, an observation that has led to significant speculation as to what this pressure may have been. One hypothesis is the presence of intermittent famines during this periods caused by crop failure, during which the ability to digest alternative energy sources such as lactose would have been more advantageous (Shennan et al., 2013; Sverrisdóttir et al., 2014). Another hypothesis is that the increased population density during this time period would have caused a larger pathogen load, thus exposing individuals to more common bouts of illness (Loog et al., 2017). As the consumption of lactose by non-lactase persistent individuals can cause diarrhea and therefore significant fluid loss (Smith et al., 2008), it is theorized that this would result in non-lactase persistent individuals having a higher mortality when exposed to these frequent illnesses. While we do not independently verify any of these hypotheses, our finding of an increase in selective pressure for lactase persistence that significantly postdates the first appearance of sustained lactose consumption in Europe supports previous hypotheses that a separate environmental event, rather than the isolated consumption of lactose itself, is what caused the strong selection for lactase persistence. We note that the selection on lactase persistence appears to be continuing to the present day, albeit with a slightly smaller selection coefficient than when the selective pressure first began. The ancestry-stratified analysis for this SNP shows that this general pattern of strong selection beginning approximately 6000 years ago followed by an attenuated selection signal in the most recent time period holds for all non-Anatolian ancestries. We notice that in the WHG ancestry, selection appears to disfavor the lactase persistence allele in the most recent time period, although sparse sampling of ancient WHG haplotypes at this locus in this time period results in high uncertainty of the true allele frequency. We therefore caution against interpreting these results as indicating modern selection against lactase persistence in this ancestry, although our results do indicate that selection is weaker than when the selective sweep first began.

Our SLC45A2 analysis appears to capture the time in which the derived allele for rs35395 was under strong selection, which was roughly during the period of 2500-5000 years ago. We find that selection during this period was particularly strong 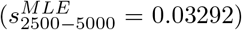, but the allele appears to not be under significant selective pressure in either younger or older time periods. We note that our finding of strong selection on depigmentation in European populations beginning 5000 years ago is in line with the results of (Wilde et al., 2014) who estimated a selection coefficient of between 2% and 10% for several skin pigmentation associated alleles. The prevailing hypothesis for the evolution of lighter skin in European populations is the resulting increased ability to synthesize vitamin D, a reaction catalyzed by solar UVB radiation, at higher latitudes where the relative concentration of solar radiation is lower (Jablonski and Chaplin, 2000). Therefore, possible explanations for the exact timing of this selective sweep include the settling of environments at higher latitudes during this period and a shift away from a hunter-gatherer diet that included vitamin D-rich fish and wild game to a vitamin D-poor agriculturalist diet. (Richards et al., 2003; Wilde et al., 2014) The ancestry-stratified analysis for this SNP shows that while all ancestries show evidence of very strong selection for skin depigmentation, the timing and strength of this selection does differ between ancestries. A possible explanation for this difference in the onset and strength of selection could be the differing latitudes inhabited by these ancestral groups through time and the possibly differing vitamin-D content of the diets of these distinct ancestral populations.

Overall, the results of our ancient DNA analysis illustrate the importance of the introduction of agriculture to Europe and the commensurate increase in population density to changing the selective pressures associated with certain traits. With the functionality of CLUES2 to test for differing selection coefficients in different time periods, it is now possible to identify these different pressures through their effect on the historic frequency of an allele.

Despite the methodological advances of CLUES2, there are nevertheless certain limitations of this approach and areas for future research. One such limitation is the effect of possible misspecification of the data-generating process. For example, we do not consider the effect of background selection, which is known to be pervasive in the human genome and to affect the observed patterns of linked neutral mutations (Pouyet et al., 2018; Johri et al., 2021; Buffalo and Kern, 2023). However, we highlight the fact that the CLUES2 framework is quite flexible, allowing for possible extensions to handle model mis-specifications simply by adjusting the relative weighting of the sampled trees in the importance sampling framework and without necessitating any changes to the ARG-inference methods.

Two possible sources of error when using CLUES2 on sequence data are improper ARG-sampling (either from poor mixing of the MCMC chain or inherent problems of the algorithms themselves) and using an insufficient number of importance samples in the Monte Carlo estimator. The degree to which these problems are an issue will depend on the exact dataset being analyzed. For example, in regions of the genome with high mutation to recombination rate ratios, ARG inference is easier and the posterior distribution of marginal gene trees will be so concentrated that few samples will be necessary (See Figure 4a). However, in datasets with low mutation to recombination rate ratios and with large numbers of samples, the state space of marginal trees may be so large and the amount of information about the length of each branch may be so low that a very large number of importance samples are necessary to effectively integrate over all the uncertainty in the data. With this in mind, to be able to more efficiently utilize large numbers of importance samples, future extensions of CLUES2 may take advantage of the fact that when evaluating the likelihood at a given selection coefficient, only a small number of samples may actually contribute substantially to the sum of the Monte Carlo estimator (Equation 1), a phenomenon known as *weight degeneracy*. (Vázquez and Míguez, 2017) This results in computational time being wasted in computing the forward algorithm for the samples which will not have a substantial contribution. Therefore, techniques from adaptive importance sampling or heuristics for choosing which subset of samples to use in the sum may be used, although this approach may encounter challenges when multiple selection coefficients are estimated. We note that when gene trees are correctly sampled and when a sufficient number of importance samples are used, CLUES2 vastly improves on summary statistic-based methods for inferring selection coefficients, which all seem to suffer from possible biases and significantly larger variances (see Figure 10). Overall CLUES2 represents a significant step forward in selection inference methodology through the use of a full-likelihood model and by being able to fully utilize the information present in large ancient DNA datasets.

## Acknowledgments

We thank the members of the Nielsen Lab at UC Berkeley for their helpful thoughts on the development of this method. In particular, we thank Diana Aguilar Gómez for her extensive testing of the software and recommendations for its input-output format and Yun Deng for his helpful discussions about HMM algorithms and associated approximations. We also thank the researchers at the Centre for GeoGenetics at the University of Copenhagen both for their feedback on CLUES2 and for hosting the authors for an extended research visit, during which much of the work for this project was done.

## A Appendix

### Differing population sizes

We also allow the specification of different population sizes in different time periods. This is accomplished by replacing *N* by a time-specific *N*_*t*_ in both the variance of the normal distribution used for the transition probabilities and the equation for the coalescence emissions. We run a set of ancient genotype simulations, identical to those described in section “Ancient Genotypes”, except that the population size in [0, 400) was 2*N* = 60000, the population size in the epoch [400, 600) was 2*N* = 200000 and the population size in the epoch [600, inf) was 2*N* = 80000. Figure 11 shows that we can properly account for changes in population size and generate accurate estimates.

**Fig 11:**
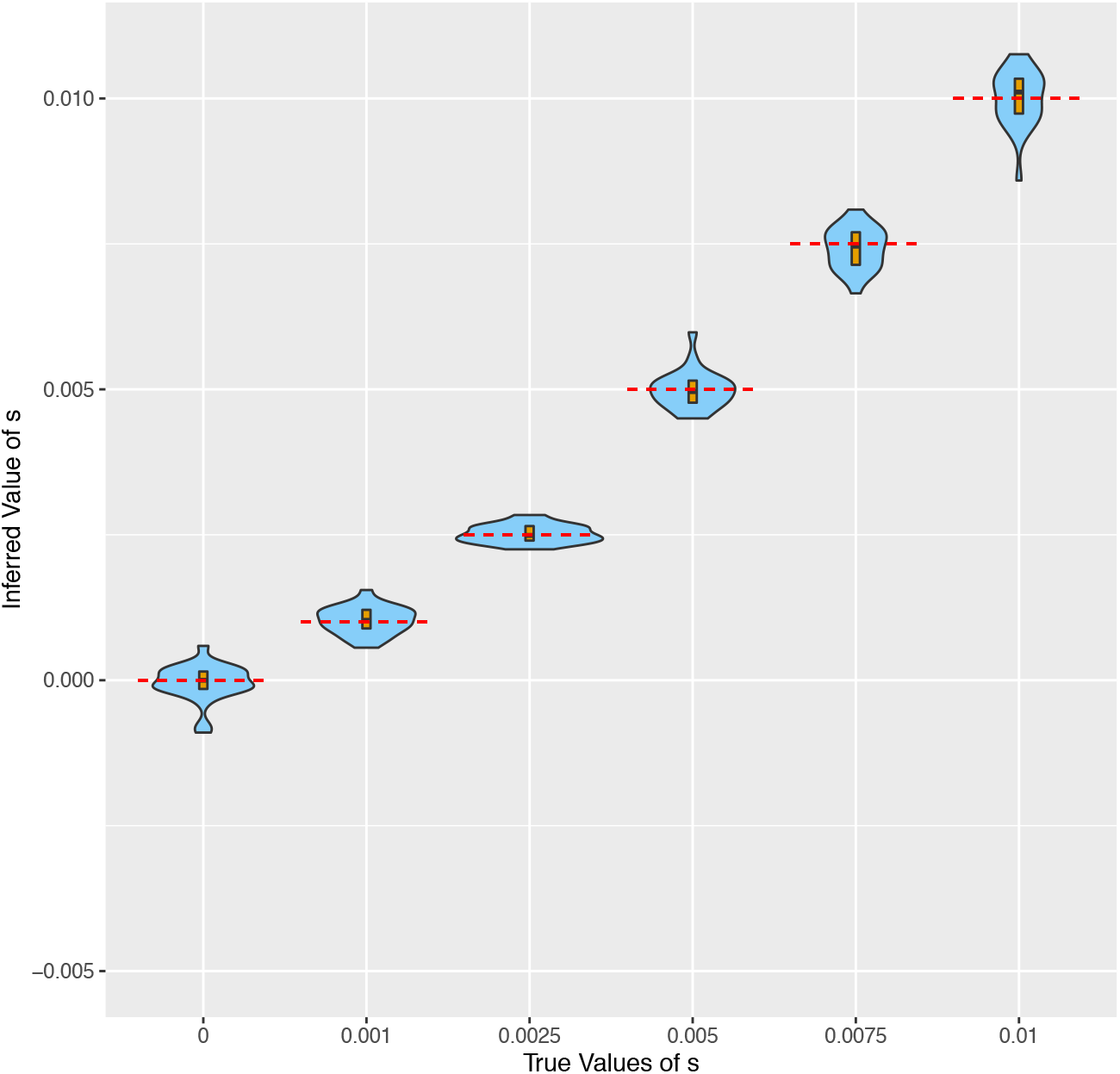
Violin plots showing the results of running CLUES2 on genotype data simulated with changing population sizes through time. Boxplots are overlayed with the whiskers omitted. True values of *s* are shown as dashed red lines. 30 replicates were performed for each selection coefficient.

### Simulation Details

#### Ancient Genotypes

We simulate ancient genotype data according to a basic forward-in-time Wright-Fisher simulation with rejection sampling. We begin our simulation with derived allele frequency 1*/N*, corresponding to a new mutation. Given the frequency at generation *k*, call it *x*_*k*_, *x*_*k*+1_ is *A*_*k*+1_*/N* where *A*_*k*+1_ is drawn from 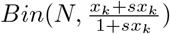. We reject simulations where the derived allele is lost and retain simulations where the derived allele is fixed. For a simulation where the allele is fixed, we then find the most recent time step *l* for which |*x*_*l*_ − *f* | *<* 0.005, where *f* is our desired modern allele frequency. We reject the simulation if no such time exists. If such a time exists, then we renumber time *l* to be the present (0 generations ago) and set the ages of the allele frequencies older than this time to be relative to this timepoint. We now have an allele frequency trajectory that goes from 1*/N* at some point in the past to *f* at the present generation. Given this trajectory, we draw one individual every *m* generations in the past (meaning one individual is sampled *m* generations in the past, one individual 2*m* generations in the past, etc) and their genotype is recorded. The exception to our approach above is the case when *s* = 0, for which we begin at time 0 with frequency *f* and simply run the Wright-Fisher process in reverse.

#### True Trees

For our simulations involving true gene trees under selection, we simulate data using the `SweepGenicSelection` function in *msprime* 1.0, which allows us to quickly and easily simulate trees under genic selection. (Baumdicker et al., 2021) As this feature is only available for strictly positive selection coefficients, our simulation results for *s* = 0 were actually simulated with *s* = 10^−6^, which makes no practical difference to the distribution of trees sampled. These trees were “recapitated” using the `recapitate` function from the pyslim library (Haller et al., 2019). in order to generate a complete, rooted gene tree. Note that the parameterization of selection in the `SweepGenicSelection` function differs from our parameterization, so we call this function using a selection coefficient of 2*s*, where *s* is the selection coefficient under which we wish to simulate.

#### Importance Sampling

For our importance sampling simulation study, we use *msprime* to simulate a gene tree under selection as described in the previous section. We then simulate neutral mutations on this tree, using a continuous genome model so that the infinite sites assumption is always satisfied. We also record the true topology of the simulated tree.

We here describe our simple MCMC algorithm. We do not vary the topology of the tree but instead use the true simulated topology. We consider this acceptable for two reasons. The first is that CLUES2 depends only on the coalescence times between allelic classes, resulting in the exchangeability of lineages within allelic classes. Secondly, the lack of recombination and our relatively small sample size (we simulate only 24 leaves) means that there will likely be at least 1 mutation per branch, resulting in only 1 possible topology that is compatible with the infinite sites assumption. Then, given a fixed topology on *L* leaves, a tree is defined by two sets of parameters:

1. The vector (*t*_*L*_ … *t*_2_) of intercoalescence times, where *t*_*k*_ denotes the time when there are *k* lineages remaining
2. The permutation vector (*p*_1_, …*p*_*L*−1_), a subset of the set of permutations on the numbers 1 to *L* − 1, which describes the relative ordering of the coalescence events. Note that not all permutations are acceptable, as many of them violate the topological constraints of the tree. For example, a coalescence node cannot occur at an older time than its parent.

We initialize the coalescence times by sampling from the prior distribution on intercoalescence times (see below). We initialize the ordering of the coalescence events in the tree using the `topo sort` function from the *igraph* package, which generates a topological sorting of the vertices of a directed acyclic graph (Csardi and Nepusz, 2005). Our target function is:

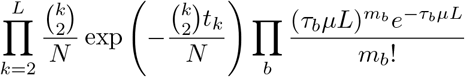

where we use *b* to index the branches of the tree, *μ* is the mutation rate, *L* is the length of the genomic region, *m*_*b*_ is the number of mutations on branch *b*, and *τ*_*b*_ is the length of branch *b*. We use a Metropolis-Hastings algorithm that is inspired by that used by Relate to sample branch lengths (see Supplementary Section 4.2 of (Speidel et al., 2019)) but differs in some minor details. With probability 1/2, we propose a change to the coalescence time vector. We pick a coalescence time *t*_*k*_ uniformly at random and sample a new time from the exponential distribution with mean *t*_*k*_. With probability 1/2, we choose a pair of adjacent indices in the permutation ordering and propose a switch between them. We automatically reject proposals that violate the topological restraints of the tree. Acceptance probabilities are calculated using the standard Metropolis-Hastings algorithm with our target function. This algorithm, while not efficient, is able to obtain samples from the distribution of branch lengths in a tree with a fixed topology and no recombination events without suffering from the observed problems with existing ARG-inference methods.

#### Gene Trees on Ancient Data

Simulating and recording full gene trees on ancient data conditional on a specified selection coefficient and a specified modern allele frequency is a challenging computational problem. However, we take advantage of the fact that the CLUES2 model implies exchangeability of lineages within each allele class to greatly simplify the simulation process. We begin by starting at the present with our specified modern allele frequency, *f*. Then, given our allele frequency at time *t* before the present, call it *x*_*t*_, we draw the derived allele frequency at generation *t* + 1 before the present as a variate from a normal distribution with mean *x*_*t*_ − *sx*_*t*_(1− *x*_*t*_) and variance *x*_*t*_(1− *x*_*t*_)*/*(2*N*). We continue this process until we reach *T*, our pre-specified time cutoff. Then, we convert the frequency at each generation into an integer number of individuals by multiplying by *N* and rounding to the nearest integer. We then initialize our *f S* derived lineages and our (1 − *f*)*S* ancestral lineages where *S* denotes the sample size of modern lineages. Going back in time, we then draw the parent of each derived lineage from the possible derived individuals in the previous generation. We do the same for the ancestral lineages. If 2 lineages have the same parent, then a coalescence is recorded for the appropriate allelic class. We continue this process until time *T*. In this way, we obtain a set of derived and ancestral coalescence times, as CLUES2 requires. It is straightforward to incorporate ancient data into this simulation procedure by specifying sampling times *τ*_1_, …, *τ*_*n*_, not necessarily unique, and adding an additional derived lineage at time *τ*_*i*_ with probability 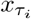 and adding an additional ancestral lineage otherwise. In this way, we can simulate the ancient gene trees input to CLUES without having to keep track of the entire tree topology. For the simulations described in the main text, we have 40 modern haplotypes. For the ancient gene tree simulations, we add 160 leaves to the tree at each of the times 100 and 200. As a comparison, we run identical simulations where we instead sample 80 diploid genotypes at each of the times 100 and 200.

## B Supplementary Figures

**Fig S1:**
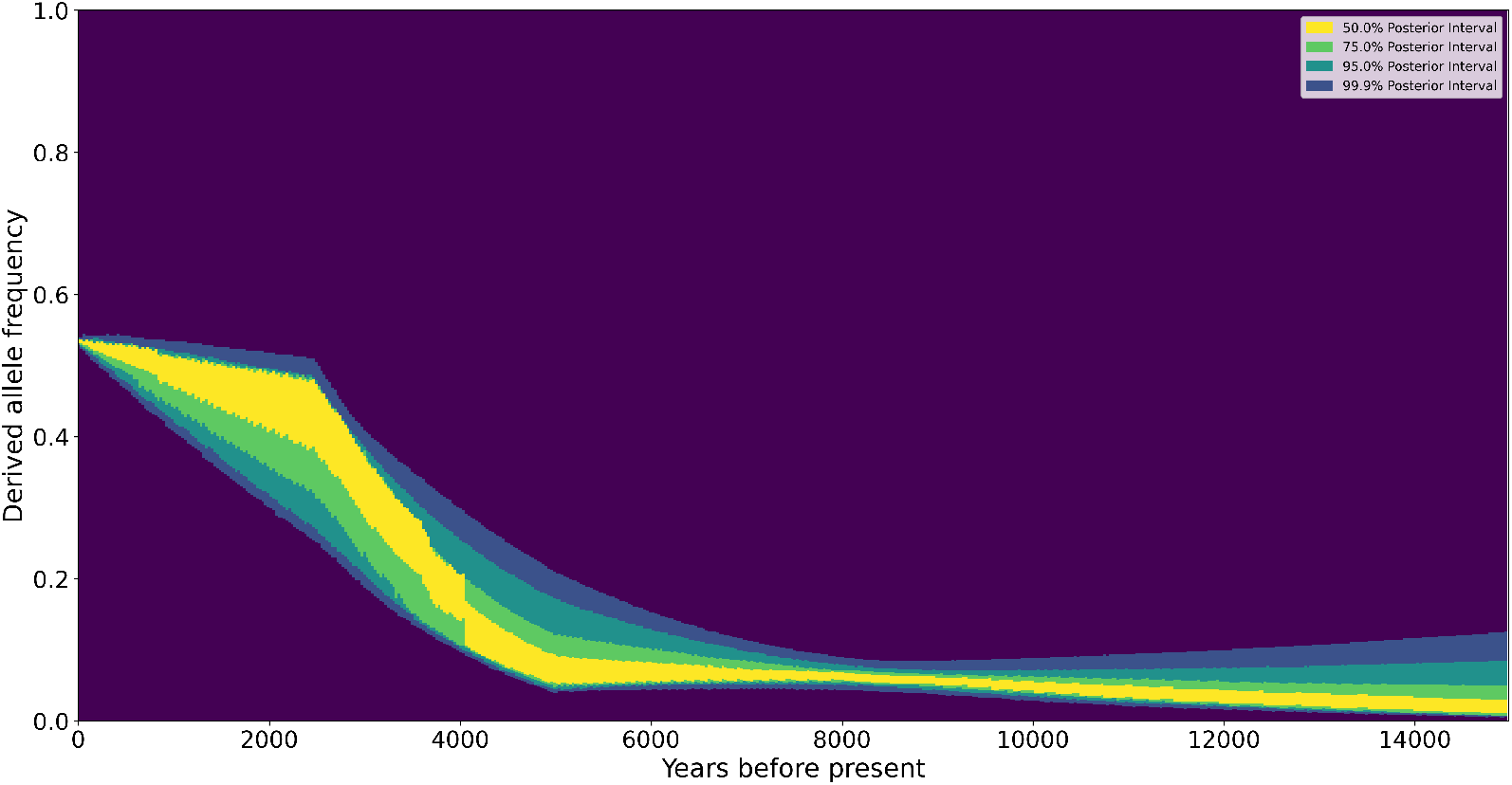
The reconstructed derived allele frequency trajectory for SNP rs4988235 in the MCM6 gene under the 3-coefficient model.

**Fig S2:**
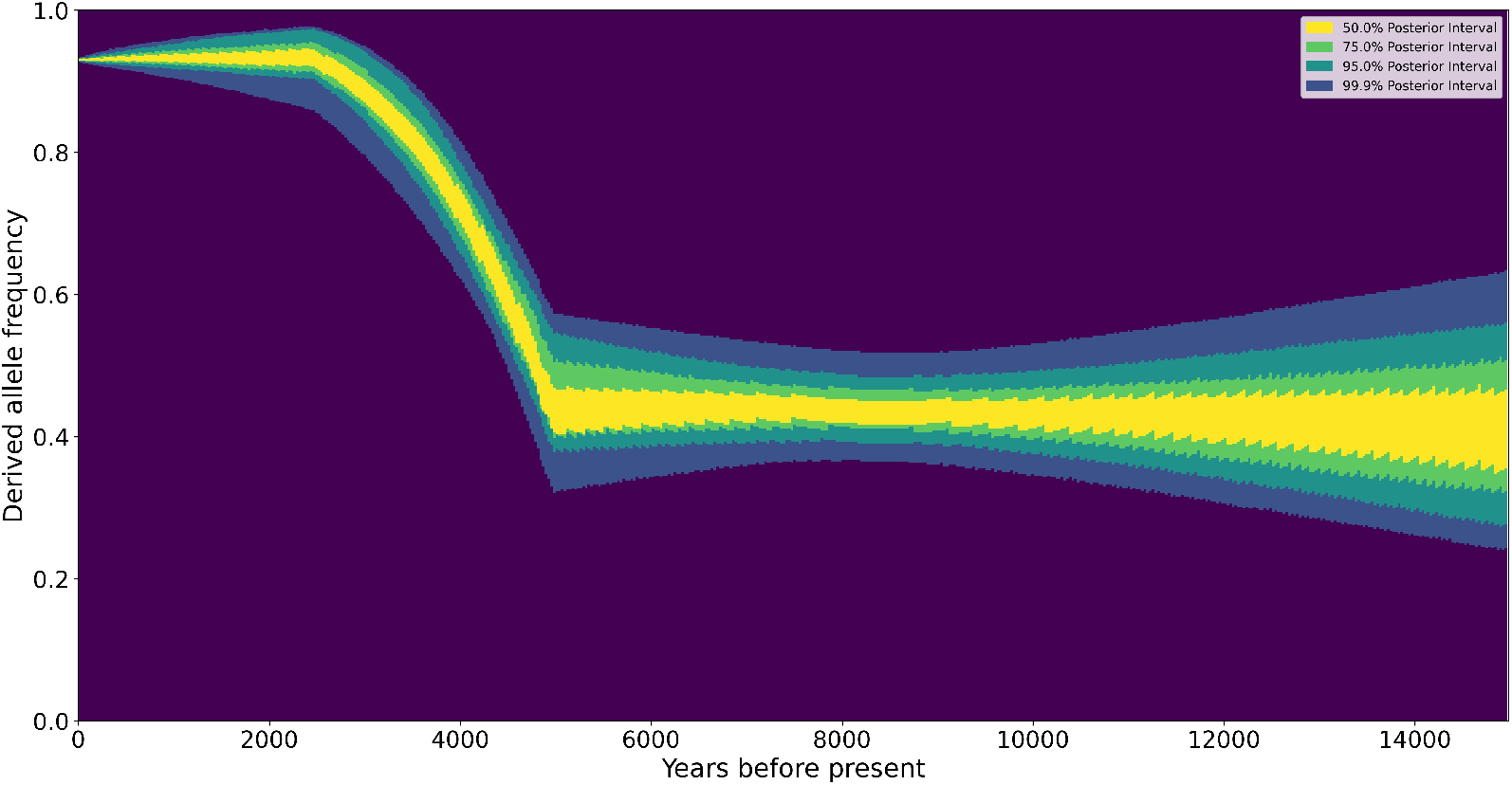
The reconstructed derived allele frequency trajectory for SNP rs35395 in the SLC45A2 gene under the 3-coefficient model.

**Fig S3:**
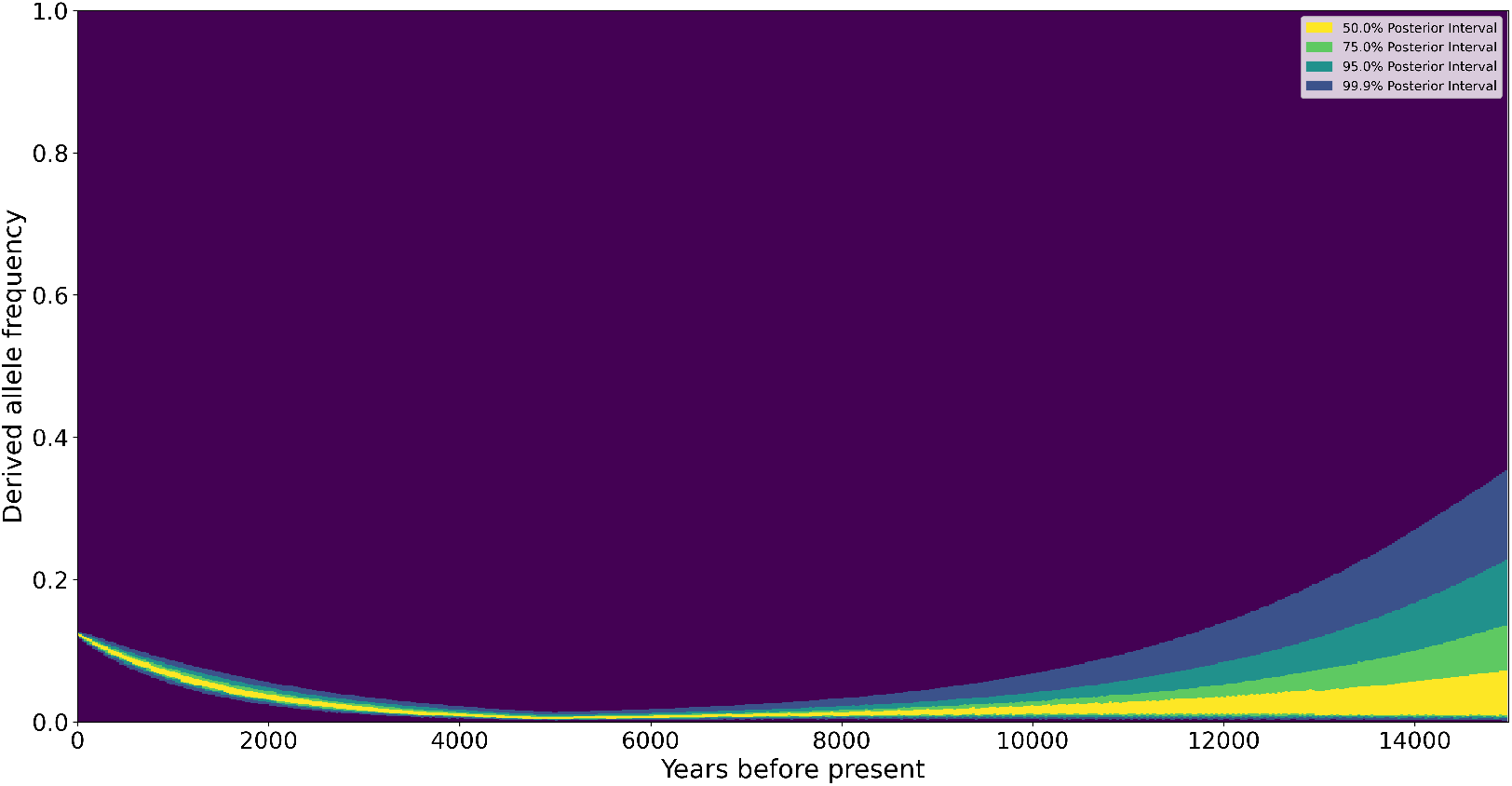
The reconstructed derived allele frequency trajectory for SNP rs12153855 in the TNXB gene under the 2-coefficient model.

**Fig S4:**
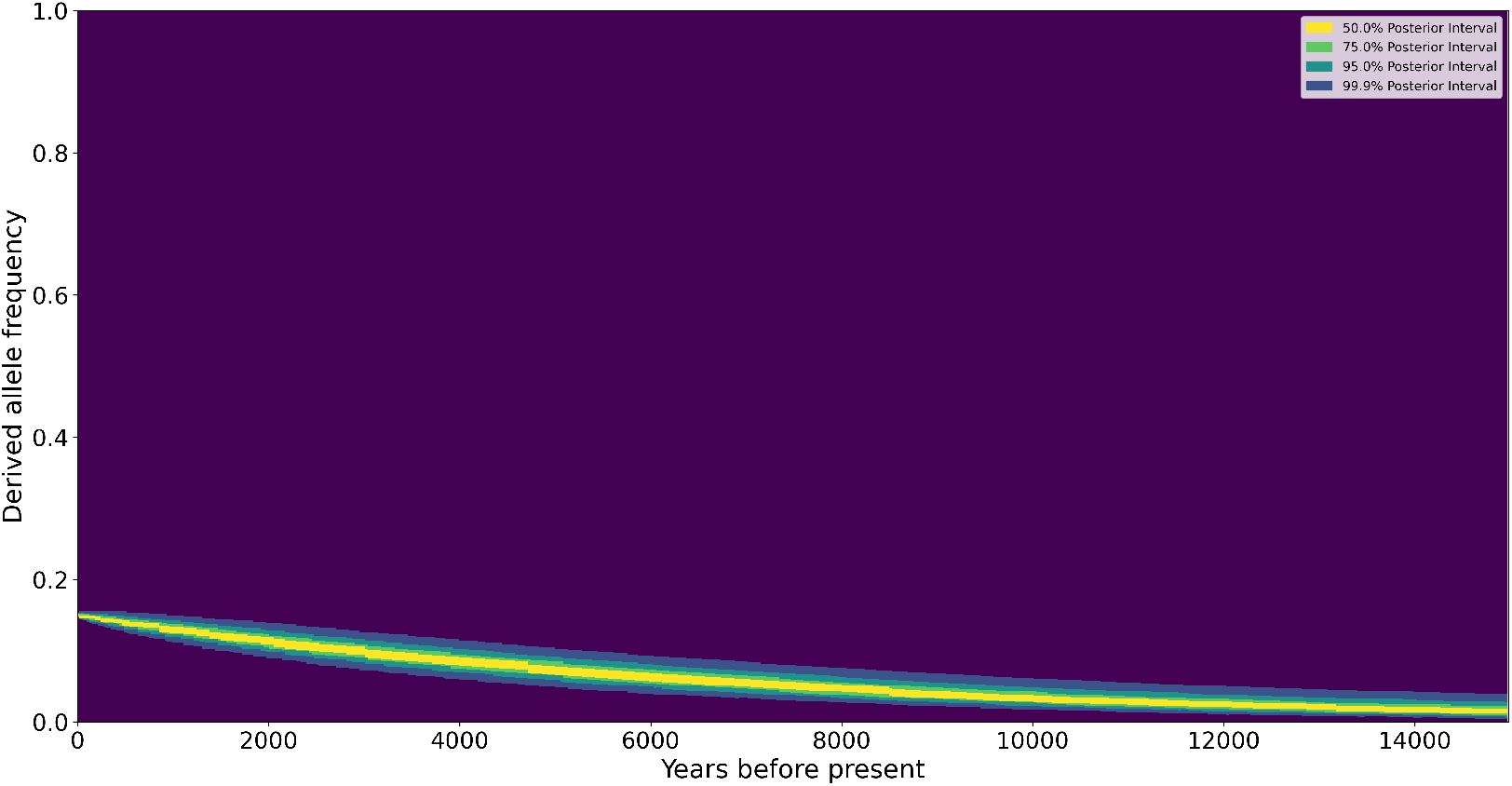
The reconstructed derived allele frequency trajectory for SNP rs75393320 in the ACP2 gene under the 1-coefficient model.

**Fig S5:**
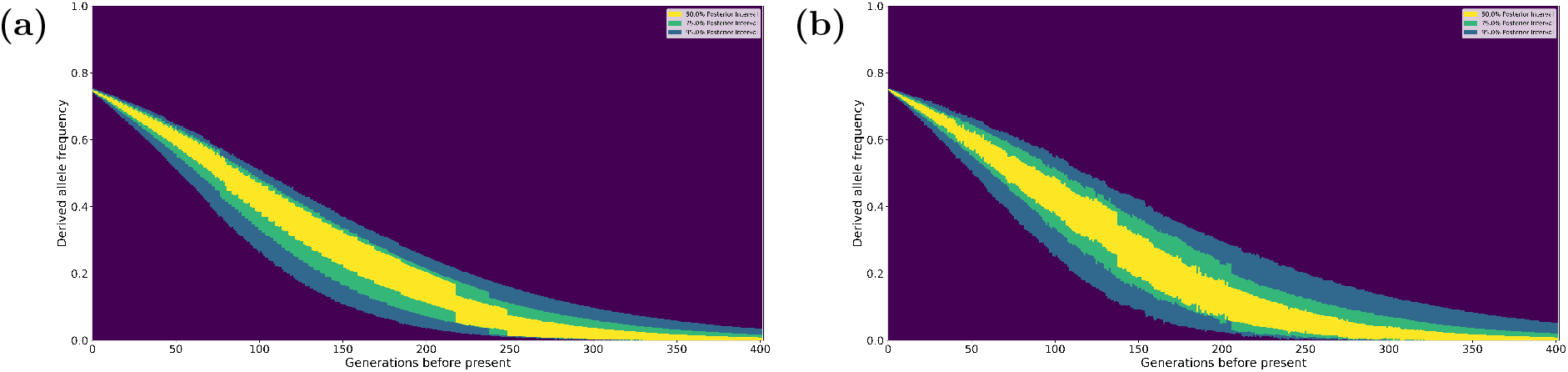
The results of the allele frequency trajectory reconstruction of CLUES2 with (a) the Monte Carlo integration framework and with (b) the exact rejection sampling approach for a 1-selection coefficient model.

**Fig S6:**
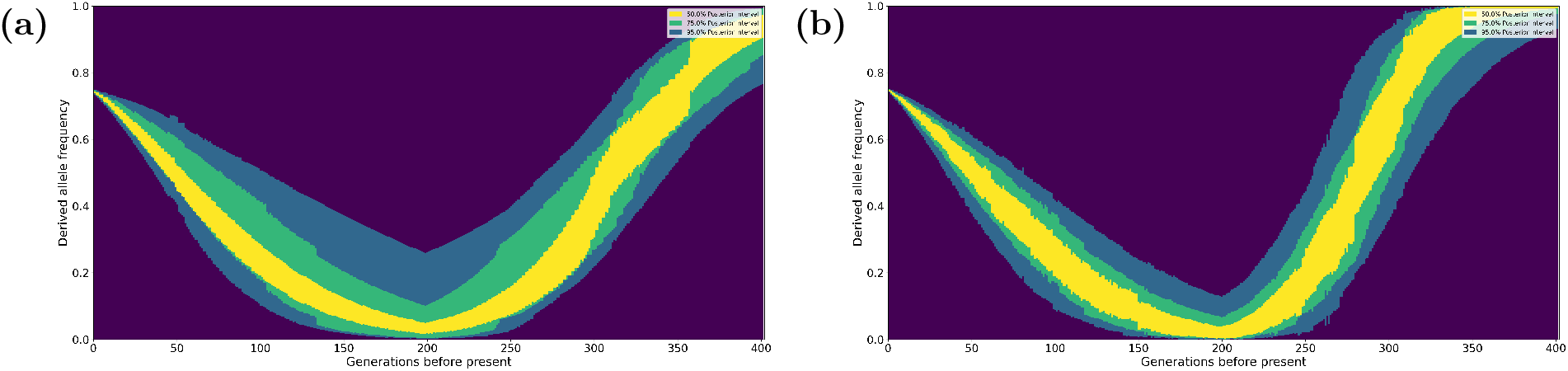
The results of the allele frequency trajectory reconstruction of CLUES2 with (a) the Monte Carlo integration framework and with (b) the exact rejection sampling approach for a 2-selection coefficient model.

**Fig S7:**
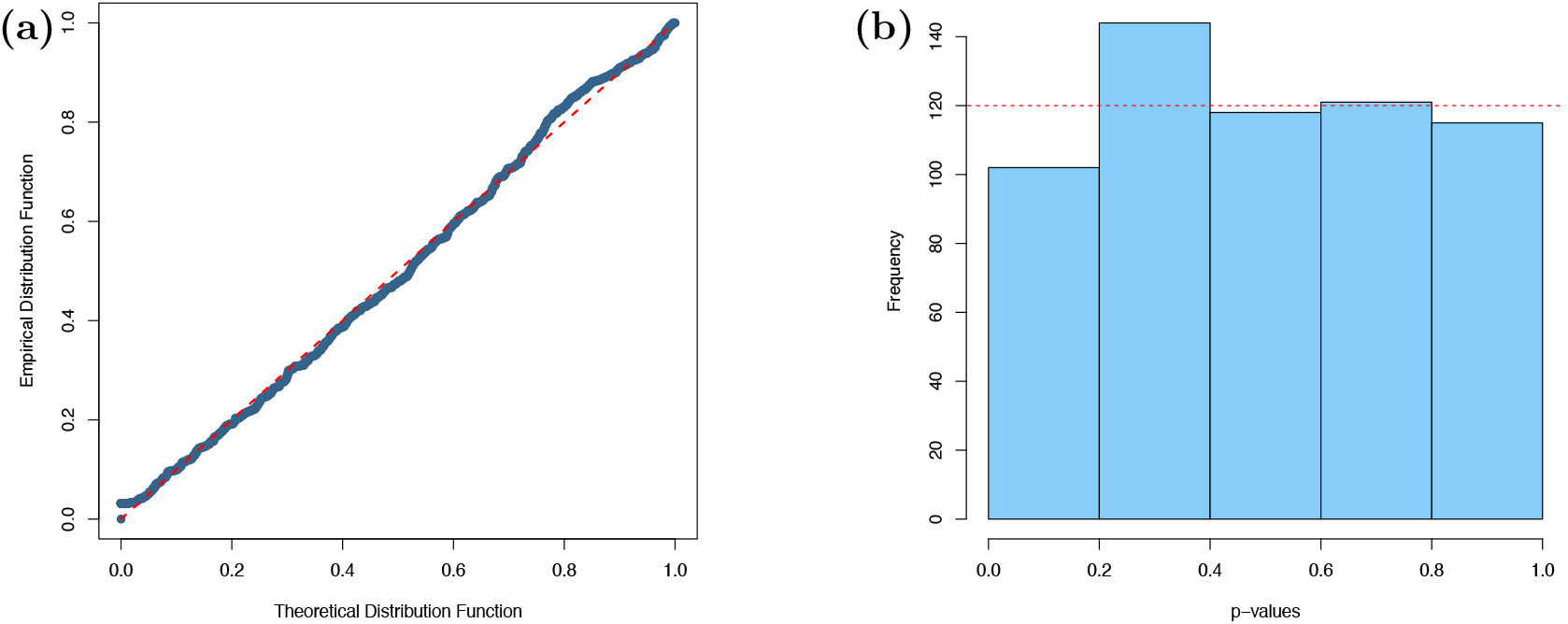
The (a) P-P plot and (b) p-value histogram for our ancient genotype simulations.

**Fig S8:**
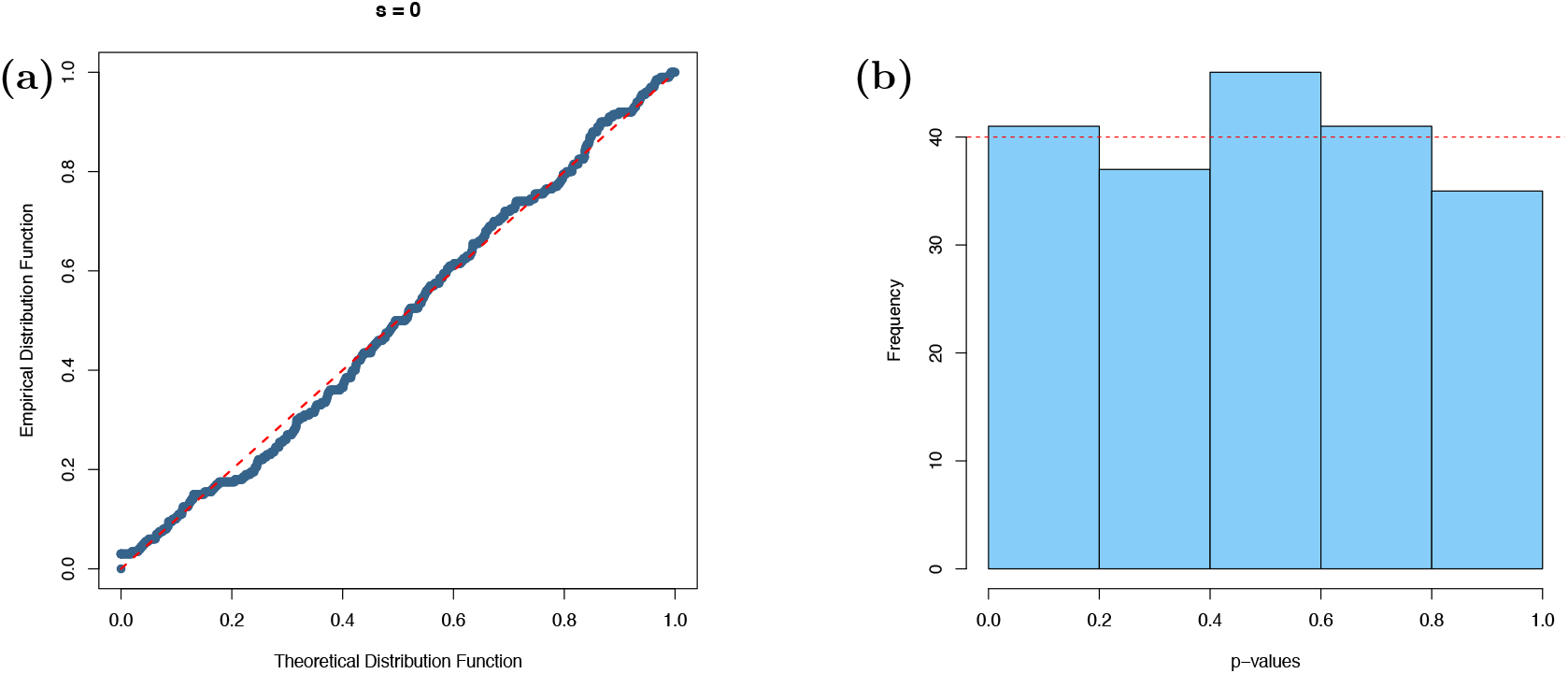
The (a) P-P plot and (b) p-value histogram for our true tree simulations.

**Fig S9:**
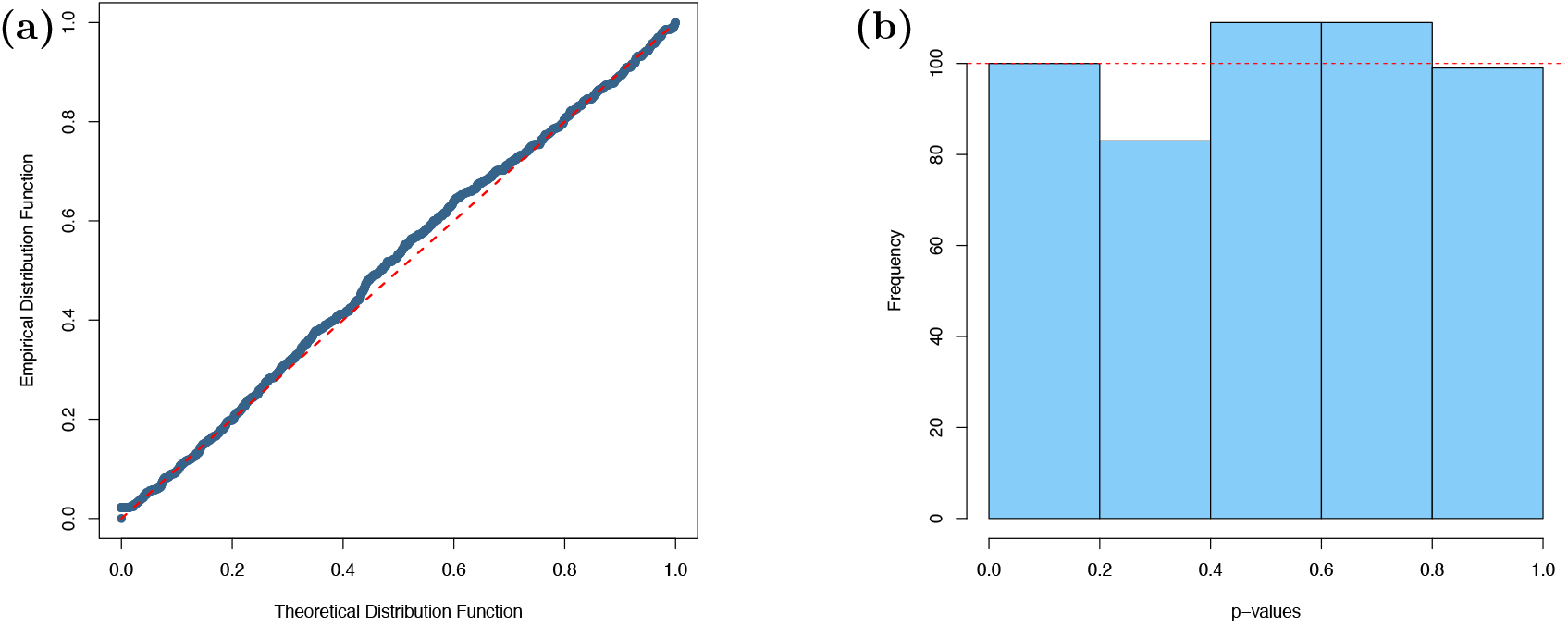
The (a) P-P plot and (b) p-value histogram for our importance sampling simulations.

**Fig S10:**
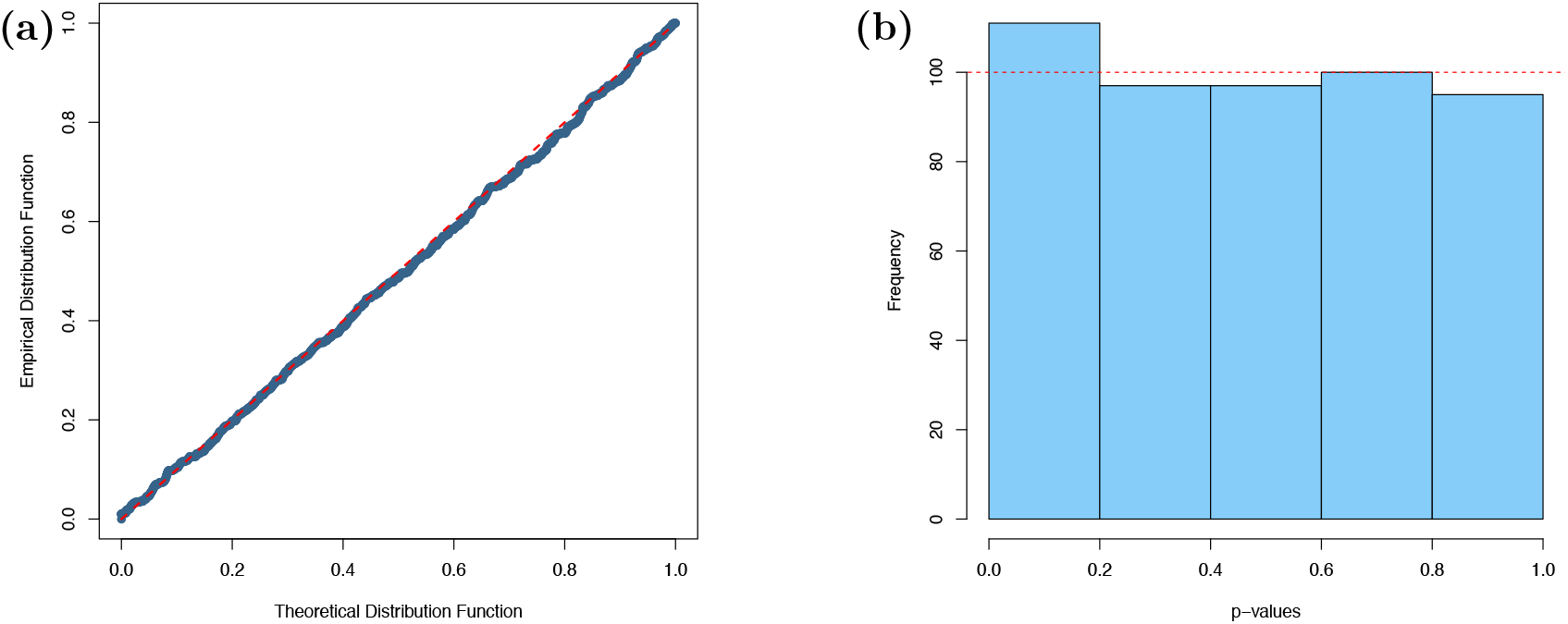
The (a) P-P plot and (b) p-value histogram for our simulations of multiple selection coefficients.

**Fig S11:**
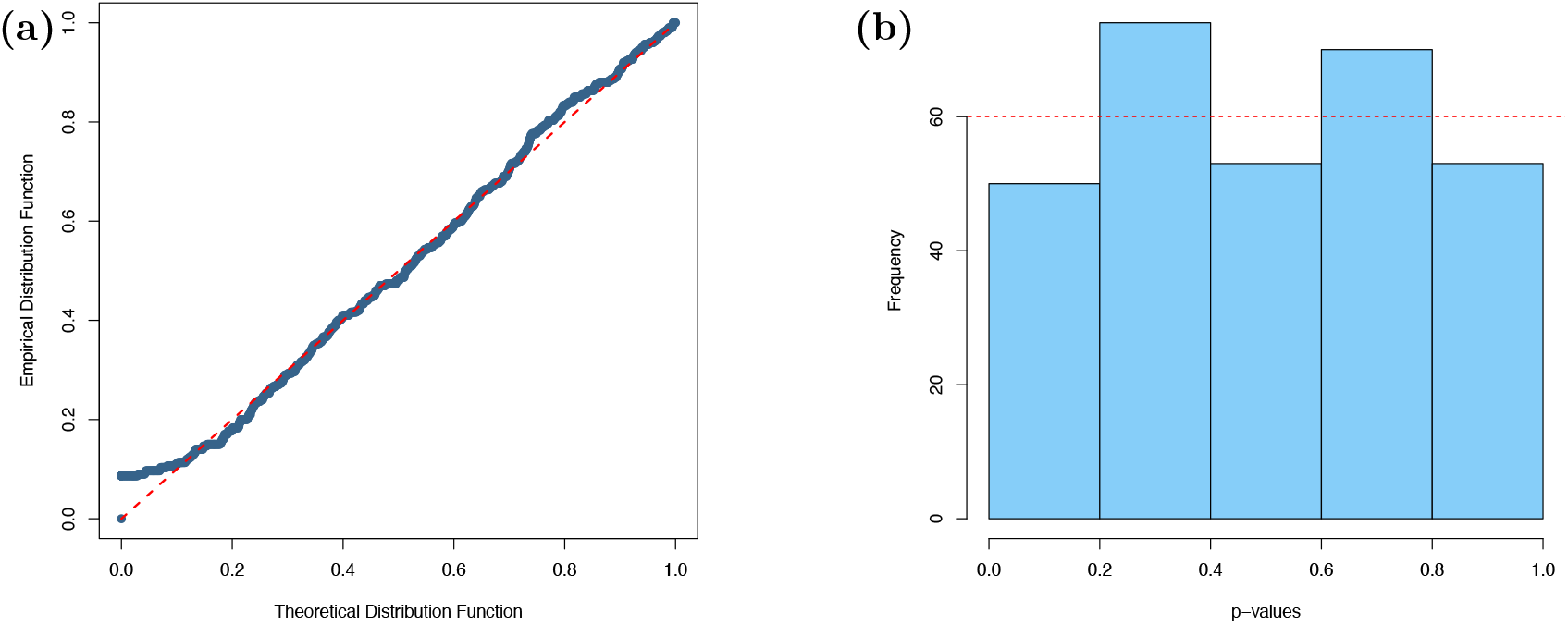
The (a) P-P plot and (b) p-value histogram for our simulations with varying population sizes.

**Fig S12:**
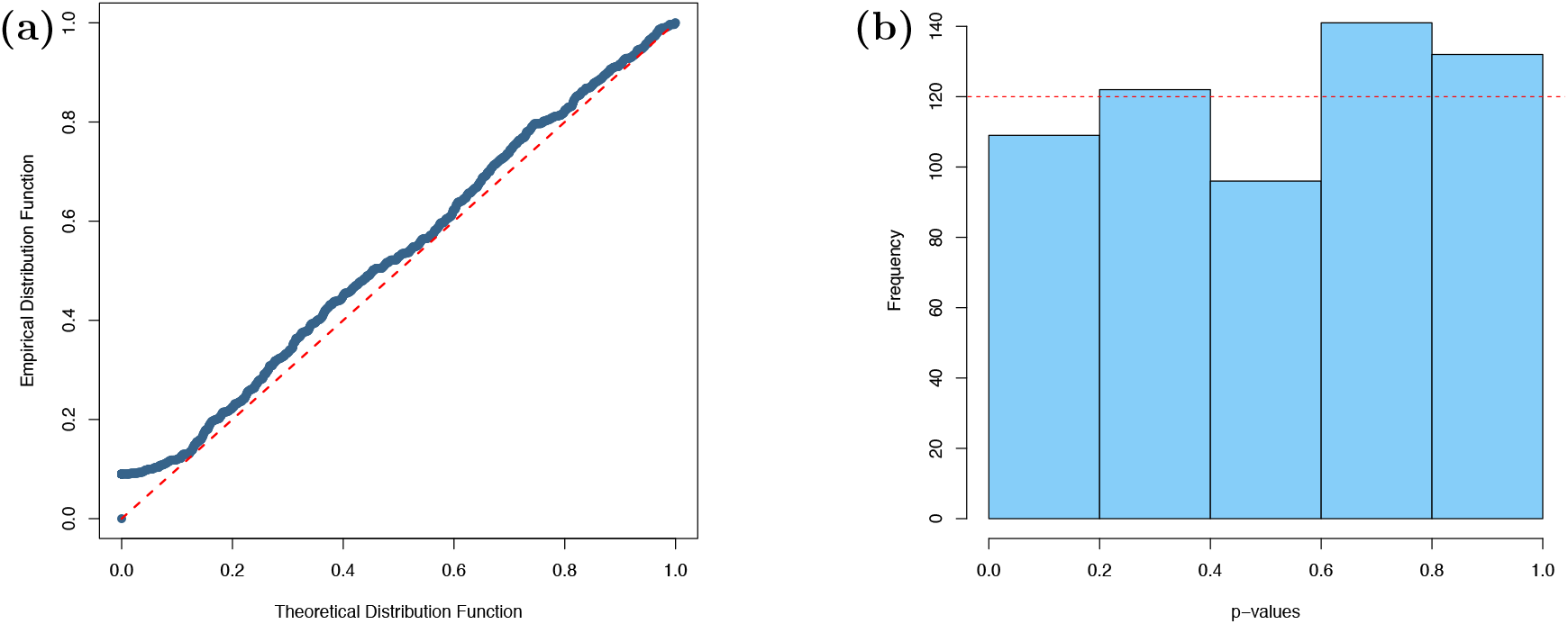
The (a) P-P plot and (b) p-value histogram for our simulations with ancient gene trees.

**Fig S13:**
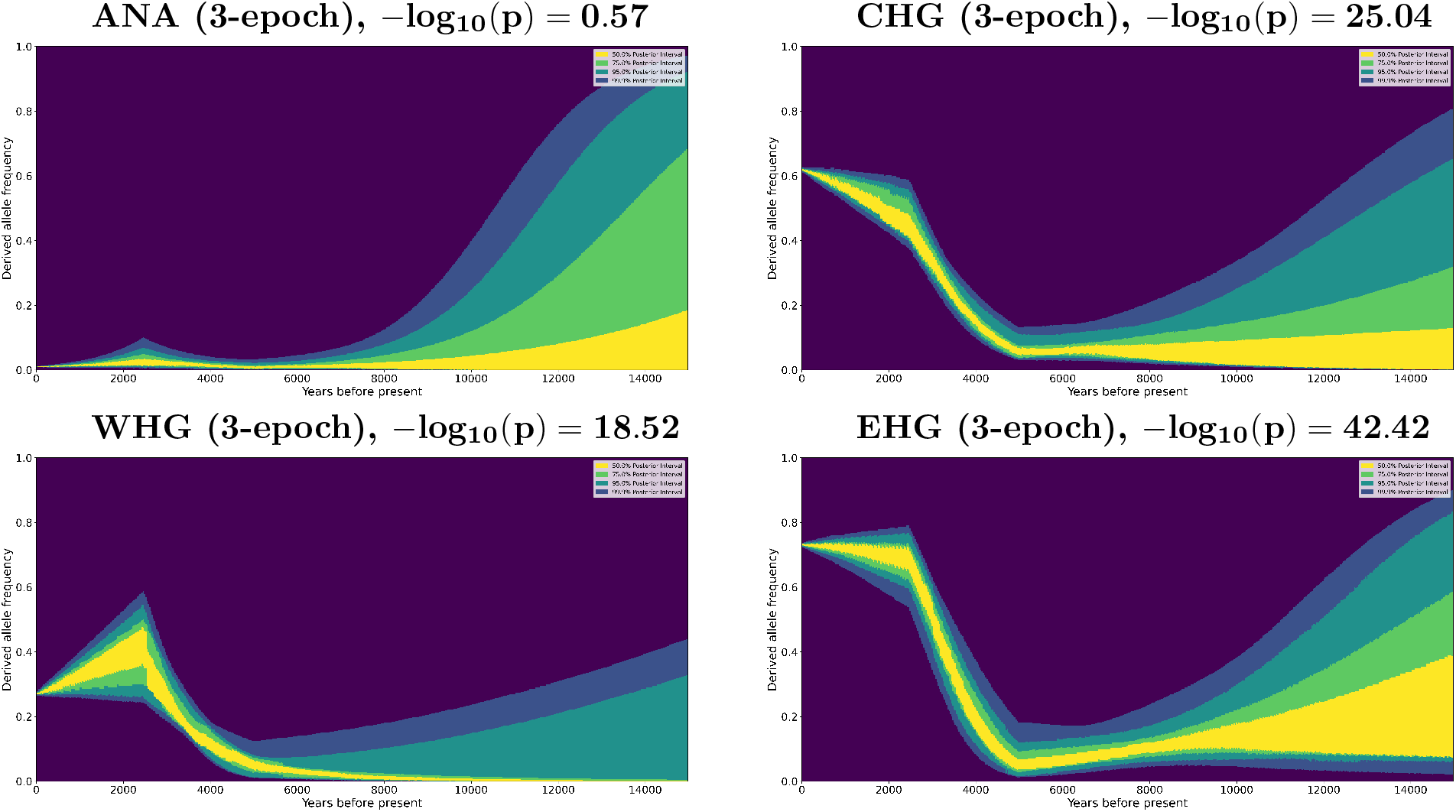
Ancestry-stratified allele frequency trajectories for the MCM6 SNP rs4988235, (chr2:136608646). Ancestries analyzed are Anatolian farmer (ANA), Caucasus hunter-gatherer (CHG), Western hunter-gatherer (WHG) and Eastern hunter gatherer (EHG).

**Fig S14:**
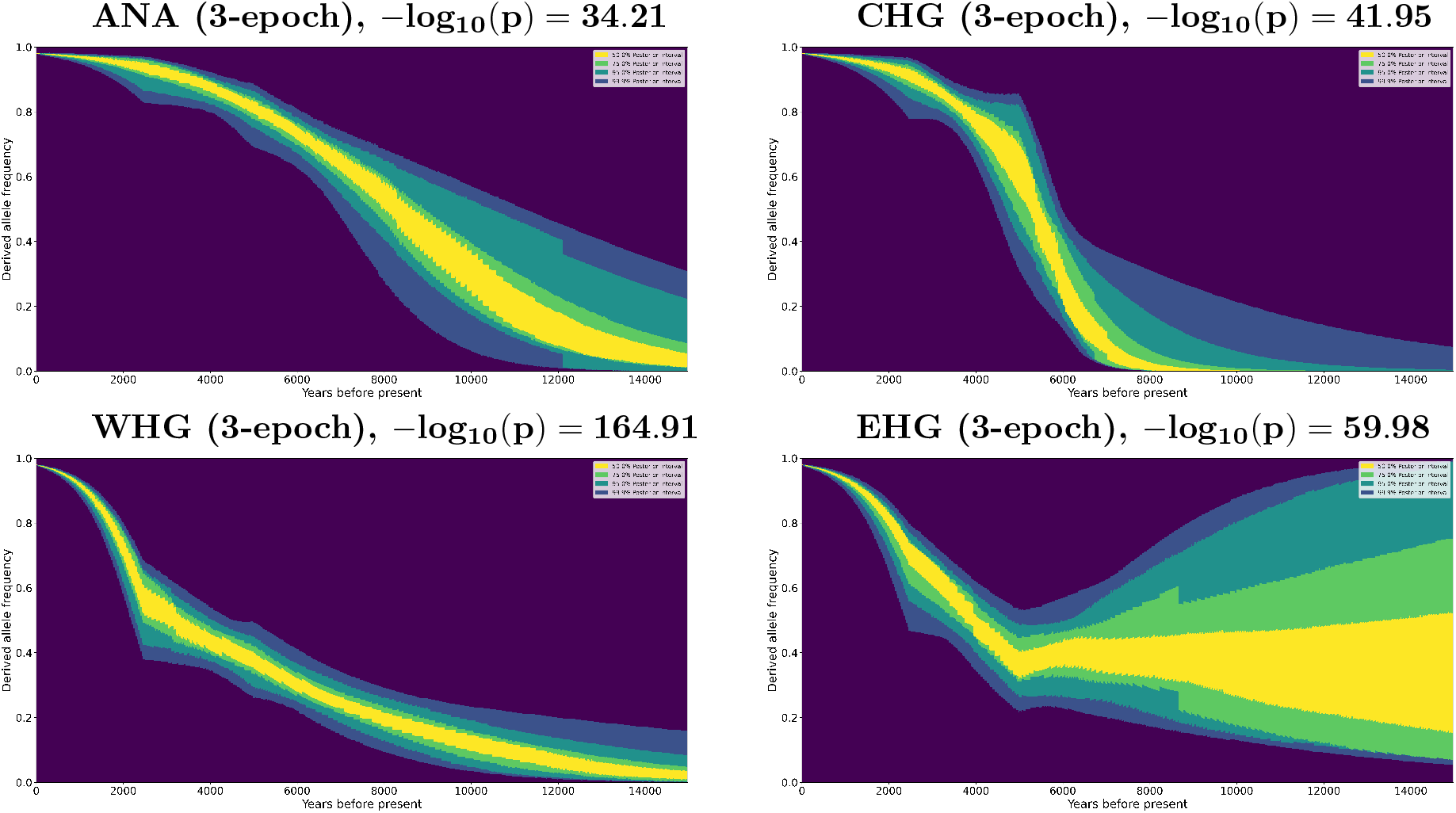
Ancestry-stratified allele frequency trajectories for the SLC45A2 SNP rs35395, (chr5:33948589). Ancestries analyzed are Anatolian farmer (ANA), Caucasus hunter-gatherer (CHG), Western hunter-gatherer (WHG) and Eastern hunter gatherer (EHG).

**Fig S15:**
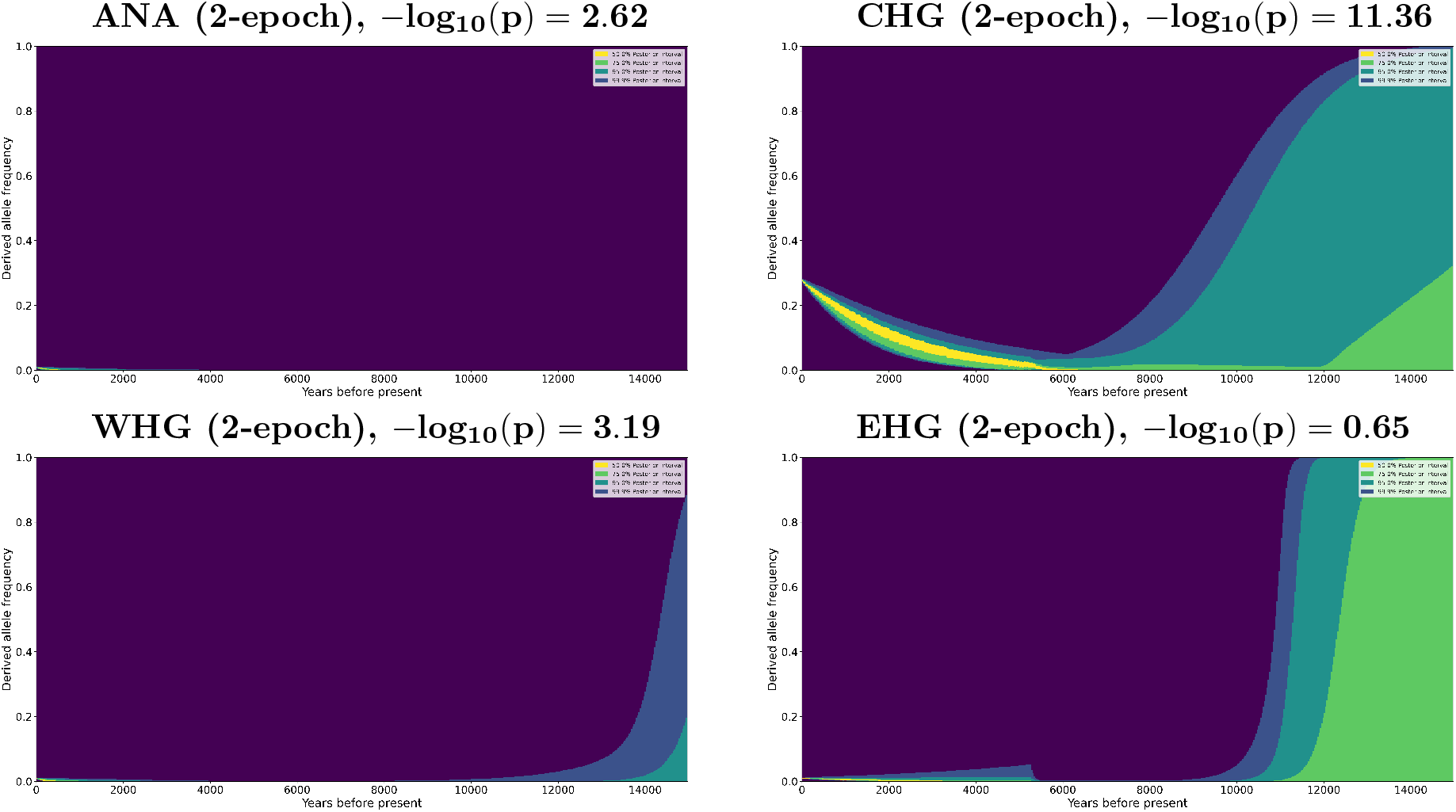
Ancestry-stratified allele frequency trajectories for the TNXB SNP rs12153855, (chr6:32074804). Ancestries analyzed are Anatolian farmer (ANA), Caucasus hunter-gatherer (CHG), Western hunter-gatherer (WHG) and Eastern hunter gatherer (EHG).

**Fig S16:**
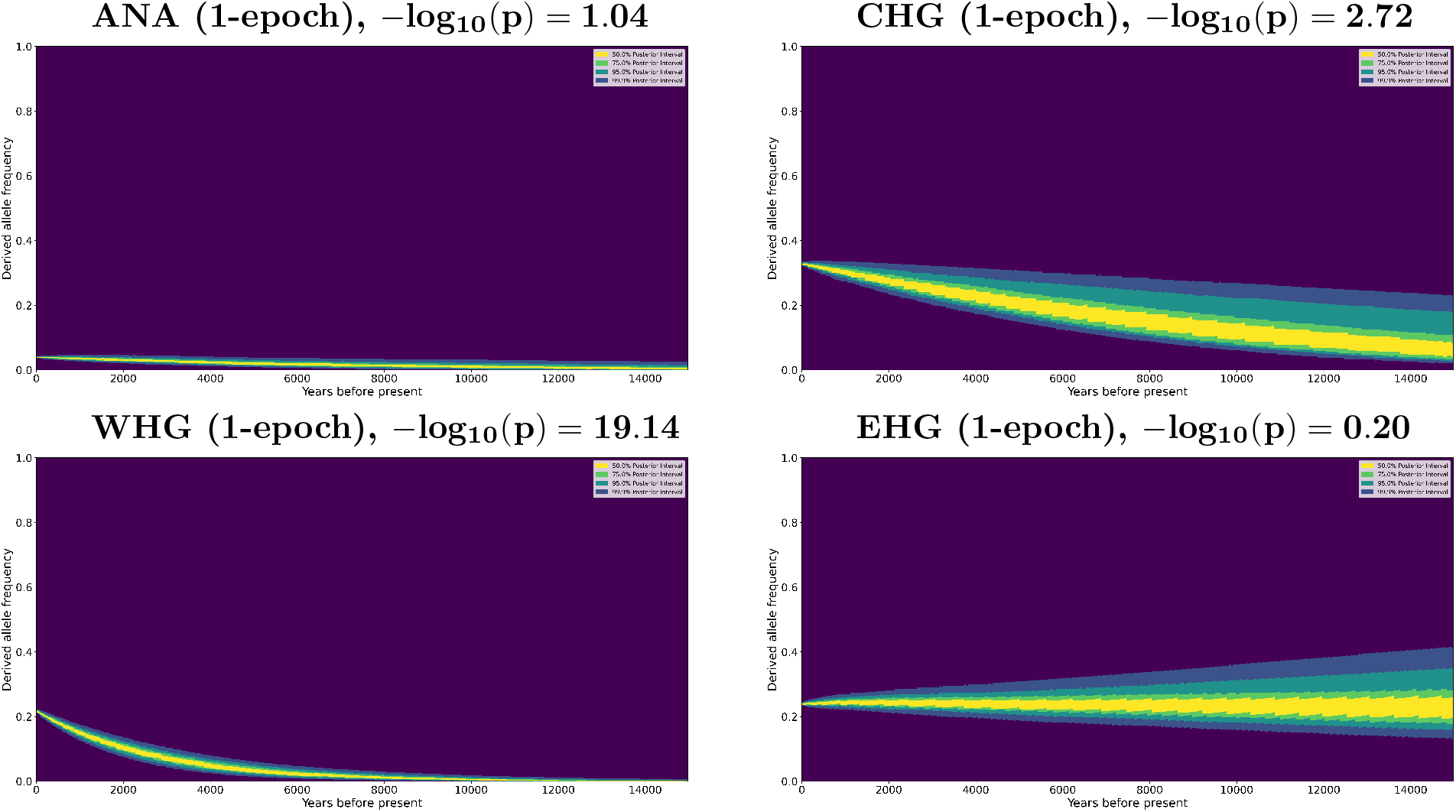
Ancestry-stratified allele frequency trajectories for the ACP2 SNP rs75393320, (chr11:47266471). Ancestries analyzed are Anatolian farmer (ANA), Caucasus hunter-gatherer (CHG), Western hunter-gatherer (WHG) and Eastern hunter gatherer (EHG).

## Notes

### Competing Interest Statement

The authors have declared no competing interest.

